# Longitudinal quantitative whole-brain microscopy reveals distinct temporal and spatial efficacies of anti-Aβ therapies

**DOI:** 10.1101/2021.01.15.426090

**Authors:** Daniel Kirschenbaum, Ehsan Dadgar-Kiani, Francesca Catto, Fabian F. Voigt, Chiara Trevisan, Oliver Bichsel, Hamid Shirani, K. Peter R. Nilsson, Karl Joachim Frontzek, Paolo Paganetti, Fritjof Helmchen, Jin Hyung Lee, Adriano Aguzzi

## Abstract

Many efforts targeting amyloid-β (Aβ) plaques for the treatment of Alzheimer’s Disease thus far have resulted in failures during clinical trials. Regional and temporal heterogeneity of efficacy and dependence on plaque maturity may have contributed to these disappointing outcomes. In this study, we mapped the regional and temporal specificity of various anti-Aβ treatments through high-resolution light-sheet imaging of electrophoretically-cleared brains. We assessed the effect on amyloid plaque formation and growth in Thy1-APP/PS1 mice subjected to β-secretase inhibitors, polythiophenes, or anti-Aβ antibodies. Each treatment showed unique spatiotemporal Aβ clearance, with polythiophenes emerging as a potent anti-Aβ compound. Furthermore, aligning with a spatial-transcriptomic atlas revealed transcripts that correlate with the efficacy of each Aβ therapy. As observed in this study, there is a striking dependence of specific treatments on the location and maturity of Aβ plaques. This may also contribute to the clinical trial failures of Aβ-therapies, suggesting that combinatorial regimens may be significantly more effective in clearing amyloid deposition.

## Introduction

Pathological protein aggregations typically occur in distinct neuroanatomical locations and give rise to specific clinical pictures (Lau, So et al. 2020, Shahnawaz, Mukherjee et al. 2020), yet the determinants of this specificity are poorly understood. In Alzheimer’s Disease (AD), the most prevalent neurodegenerative disorder (Fiest, Roberts et al. 2016), deposition of amyloid-β (Aβ) plaques occurs in a well-characterized sequence (Braak, Braak et al. 1993, Hardy and Selkoe 2002, Thal, Rub et al. 2002). This plaque load can be effectively reduced by either quenching Aβ production(De Strooper, Vassar et al. 2010), reducing the propagation of Aβ aggregates (Jiang, Liu et al. 2013), or enhancing Aβ catabolism (Sevigny, Chiao et al. 2016). However, the clinical efficacy of Aβ removal is still debated (Morris, Clark et al. 2018, Howard and Liu 2020), perhaps because intervention is too late to be efficacious (Sperling, Karlawish et al. 2013). It is also conceivable that anti-Aβ drugs remove plaques differentially in distinct CNS regions, some of which may not coincide with the areas that matter most to proper brain functioning.

To investigate this latter hypothesis, we developed a high-throughput quantitative 3D histology (Q3D) platform for optically clarifying, staining, imaging and quantifying Aβ plaques in whole brains of mice. Aβ plaques were electrophoretically stained in cleared brains and imaged with a mesoscale selective plane illumination microscope (mesoSPIM) (Voigt, Kirschenbaum et al. 2019). In APP/PS1 mice(Radde, Bolmont et al. 2006), we tested the effects of the polythiophene LIN5044, which intercalates with amyloids and is therapeutic in prion diseases (Margalith, Suter et al. 2012, Herrmann, Schutz et al. 2015), the BACE1 inhibitor NB360 (Neumann, Rueeger et al. 2015, Neumann, Machauer et al. 2019), and a β1-antibody (Paganetti and Schmitz 1996). We found that each drug had differential efficacy on plaque formation, plaque growth, and plaque maturity, as well as a striking spatiotemporal dependence. We further found that the two most effective treatments for reducing plaque growth, BACE1 and LIN5044, acted onto distinct, largely non-overlapping brain regions. Finally, the alignment of whole-brain treatment maps to a spatial transcriptomics atlas allowed us to identify transcriptional signatures correlating with the effectiveness of each drug.

## Results

### Rapid tissue clearing and staining platform

Detergent-mediated lipid extraction from hydrogel-embedded tissues is facilitated by electrophoretic mobilization of detergent molecules (Chung, Wallace et al. 2013, Tomer, Ye et al. 2014) in a buffer-filled container. However, the electrical resistivity of 4% paraformaldehyde-fixed PBS-soaked brain tissue is 4-fold higher than that of PBS (**Figure S1E**). Therefore, any buffer surrounding the specimen short-circuits its electrophoresis. We resolved this issue by constructing a focused electrophoretic clearing (FEC) device that uncouples buffer recirculation in the anodic and cathodic circuits with an insulating layer (**Figure 1A and S1**). By forcing the electrical current to traverse the tissue specimen (130 mA in constant-current, 39.5 °C), FEC reduced the clearing time from 48-120 with CLARITY to 6-14 hours (Chung, Wallace et al. 2013, Tomer, Ye et al. 2014) and resulted in homogeneous high-quality clearing (**Figure 1B-C and S1G - H**). Similar to other hydrogel-based clearing methods, brain tissue showed some swelling during clearing, which was reversible upon PBS washing and refractive-index matching.

**Figure 1.**
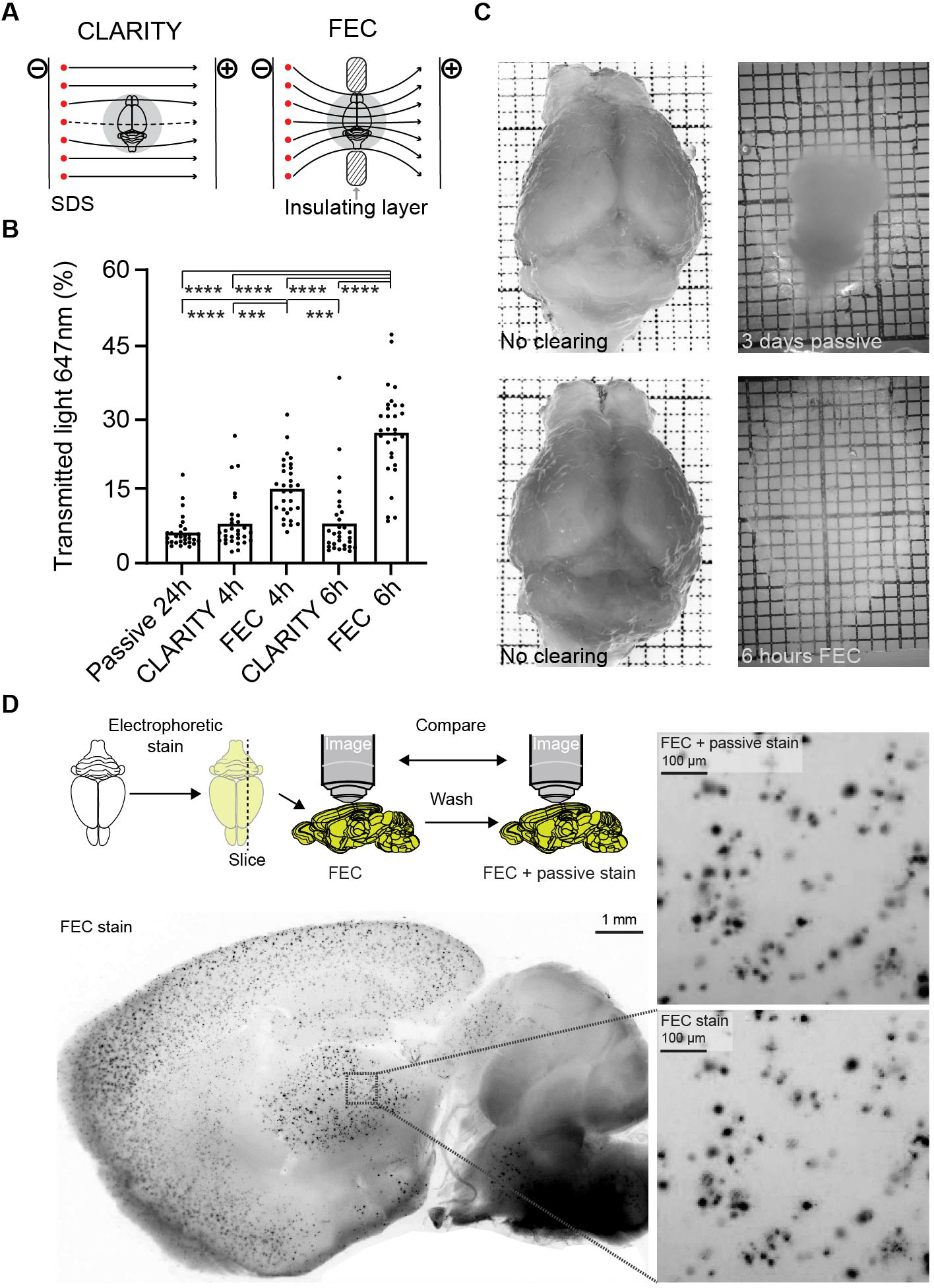
Focused electrophoresis improves tissue clearing efficiency. (**A**) Schematics of both CLARITY and focused electrophoretic clearing (FEC). An insulating layer constrains the electrical field through the tissue. (**B**) Light transmittance of tissue increased more rapidly with FEC than with CLARITY. Each datapoint plotted is the mean from three neighboring transmittance readings (hence resulting in 10 datapoints from the 30 measurements/brain) (One-way ANOVA *** p<0.001, **** p<0.0001). (**C**) Comparison of mouse brains cleared by FEC (6 hrs) and passive (3 days). Passive clearing was incomplete even after 3 days. **(D)** After whole-brain electrophoretic staining with polythiophenes, sagittal 500 µm slices were cut and plaques were counted. By passively re-staining the same slice, no additional plaques were detected (“FEC + passive”). Focal shifts after slice reprocessing account for the slight differences between the images.

For Aβ plaque staining of intact mouse brains by electrophoresis, we constructed buffer-filled chambers hosting the electrodes. Their inner faces were cast with 10% polyacrylamide in tris-tricine buffer and functioned as electrically conductive contact surfaces. The sample was mounted with a holder between the two polyacrylamide walls. This allowed electrophoresis to occur through the buffers, the gels and the tissue specimen; spanning 10 cm between the two electrodes (**Figure S2A-B**). The electric resistance of the electrophoretic system (20 V, 20 °C) increased from initially ∼2 kΩ to ∼20 kΩ after 2 hours (**Figure S2C**).

We then ran native-gel electrophoreses of proteins with various charges at different pH and ionic strengths. Tris-tricine (50 mM each) at pH 8.5 yielded the best results (**Figure S2D-G**). As expected, the electrophoretic mobility of proteins was influenced by the charge of covalently coupled fluorophores (**Figure S2D-G**). The polythiophenes qFTAA (616.5 g/mol, m/z = 205.5) and hFTAA (948.9 g/mol, m/z = 237.125) were dissolved in agarose (600 µl, congealing temperature 26-30 °C) and cast on the acrylamide-tissue interface in order to confine them to the smallest possible volume. Under these conditions, the dye front traversed the entire brain within 2 hours.

An uneven passage of the dye front through the brain may lead to local inhomogeneities of plaque detection, particularly at gel-liquid interfaces. To investigate this question, a hydrogel-embedded and cleared APP/PS1 brain was electrophoretically stained with polythiophenes (2 hours) and cut into 500-μm sagittal sections with a vibratome. Free-floating sections were imaged with a fluorescent stereomicroscope. Then, the sections were passively re-stained with the same polythiophene dyes using a well-established protocol (Nystrom, Psonka-Antonczyk et al. 2013, Rasmussen, Mahler et al. 2017) and images were acquired again. The numbers of Aβ plaques were 3085 and 3061 plaques before and after re-staining, respectively, and their morphology was very similar (**Figure 1D, S3**). Hence the sensitivity and spatial homogeneity of electrophoretic plaque staining of whole brains was not inferior to that of conventional histochemical slice staining. Slight differences in plaque counts and morphology were a result of physical distortions of the slices during passive staining, and due to focal shifts during re-imaging of the free-floating slices.

Antibody Aβ17-24 recognizes early plaques and stains their entire surface, whereas N3pE labels plaques which accumulate at later stages (Rijal Upadhaya, Kosterin et al. 2014). Likewise, the polythiophene hFTAA stains the entire area of early plaques whereas qFTAA stains the cores of plaques in older mice. The qFTAA/hFTAA ratio correlates with plaque compactness (Nystrom, Psonka-Antonczyk et al. 2013) and is used as a proxy for their maturity. We stained histological sections (3 µm) from paraffin-embedded APP/PS1 brains with Aβ17-24 or N3pE, followed by staining with qFTAA and hFTAA. The hFTAA and Aβ17-24 signals were largely superimposable and identified more plaques than qFTAA and N3pE, which stained selectively the cores of a subset of plaques (**Figure S4A**). Most Aβ17-24^+^ N3pE^-^ plaques were hFTAA^+^ qFTAA^-^, suggesting that they contained less mature amyloid. We conclude that the qFTAA/hFTAA stain is a good proxy to plaque maturity and suitable for whole brain staining (**Figure S4B - C**).

### Evaluation of anti-Aβ therapies by Q3D

Groups of 2-month old or 11-month old APP/PS1 mice (30 and 25 mice/group, henceforth referred to as “young” and “old”, respectively) were treated for 90 days with the BACE1 inhibitor NB360 (0.5 g inhibitor/kg chow, ∼3 mg inhibitor/day/mouse), with β1 antibody against Aβ (0.5 mg in 200µl PBS, 1x/week intraperitoneally, based on previous protocols (Pfeifer, Boncristiano et al. 2002, Balakrishnan, Rijal Upadhaya et al. 2015), or with the amyloid-binding compound LIN5044 (0.4 mg in 100µl PBS, 1x/week intraperitoneally) (**Table S1**). Control treatments included control food chow and intraperitoneally injected recombinant pooled IgG or PBS, respectively (**Figure 2A**). Mice were sacrificed one week after the last administration of LIN5044 or β1; the NB360 chow was provided without interruption. Brains were subjected to clearing, staining, and imaging. Raw data volumes were transformed to the coordinate space of the Allen Brain Atlas (Wang, Ding et al. 2020) and anatomically registered (**Figure S5D-E**). We then performed automated plaque segmentation and regional quantification of plaque pathology (**Figure 2B, S5A-C, Table S2**). Voxel-level plaque counts, mean size, and maturity (qFTAA/hFTAA ratio) were determined for each treatment group (**Figure 2C-D, S6, Table S3**). Corresponding voxels of brains treated with anti-Aβ compounds and their respective controls were compared pairwise by inferential statistics (**Figure 2E**). This allowed us to identify “Significantly Altered Voxels” (SAV) across entire brain volumes. SAV heatmaps were presented as montages of coronal slices.

**Figure 2.**
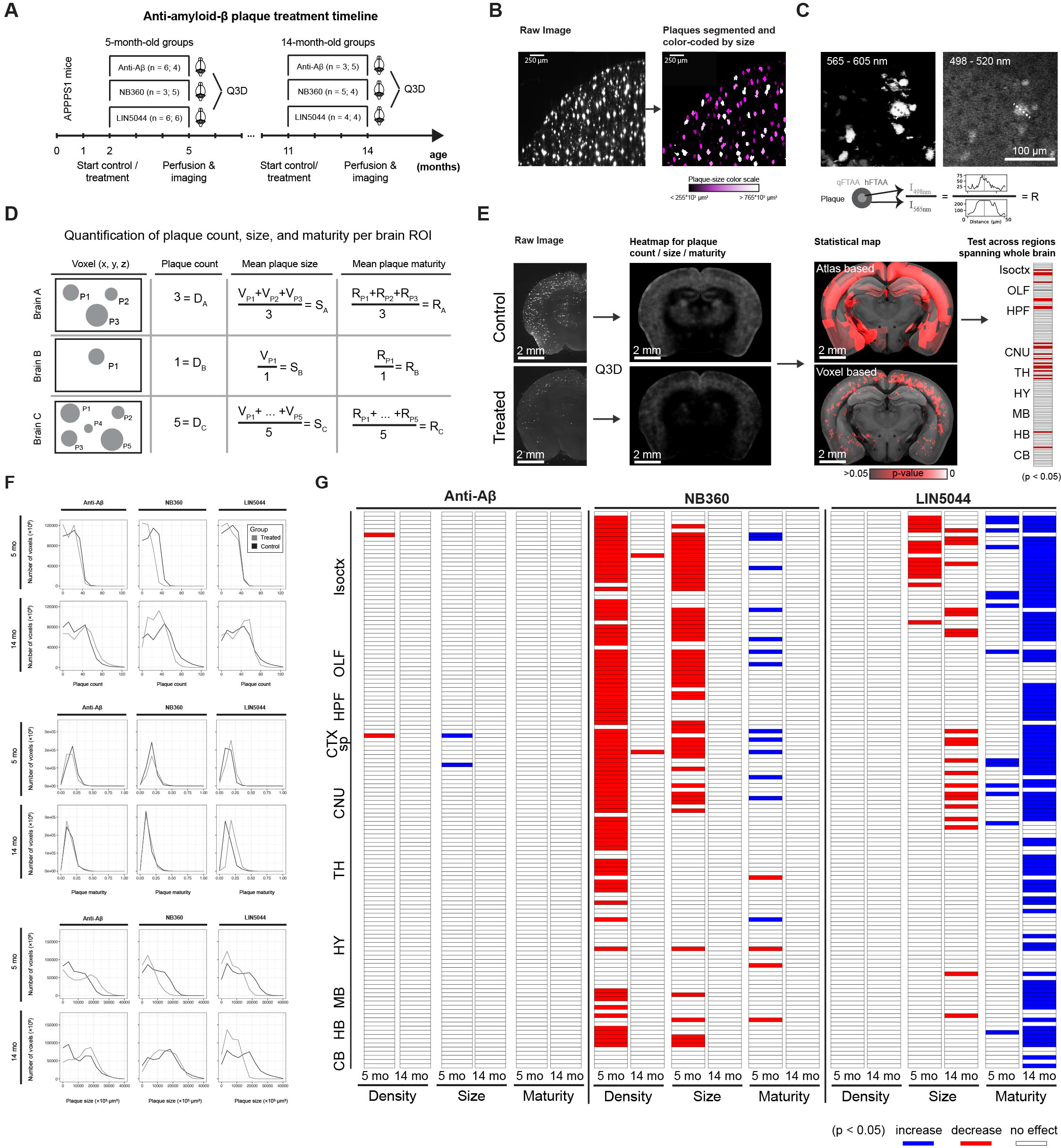
Region and age-specific plaque clearing by various anti-Aβ treatments. (**A**) APPPS1 mice were treated with antibodies, the BACE1 inhibitor NB360, the polythiophene LIN5044 or appropriate controls. (B) We segmented the plaques and color-coded them by plaque size (um^3^). (C) Intensity ratios were calculated by dividing the peak fluorescent emission in the qFTAA channel with the peak emission of the hFTAA channel in the center of each plaque. (**D**) Descriptive statistics were calculated for every voxel of individual brains resulting in plaque counts, mean plaque size and mean intensity ratio for each voxel (plaques: P1, P2 … Pn). (**E**) After every brain was registered to a reference atlas, plaques were grouped by anatomical brain regions. The mean plaque density, size and maturity was compared between control and treated brains for corresponding brain regions. Seeking an unbiased volume unit for spatial analysis we divided brain data into (25 µm)^3^ voxels. Atlas-based anatomical normalization overestimates the volume of treatment-affected brain compared to voxel-level analysis. (**F**) Histograms visualizing the changes in plaque size and maturity upon different treatments. For the β1 antibody treatment, there was a reduction in the number of smaller plaques, but there was no change in plaque density or maturity. NB360 and LIN5044 reduced the prevalence of large plaques in young mice; LIN5044 also reduced it in old mice. NB360 affected the plaque maturity of young but not of old mice, while LIN5044 increased plaque maturity primarily in old mice. (**G**) Statistical tests resulted in heatmaps of significance by anatomical brain regions. The β1 antibody treatment showed very limited effects in all analyzed brain regions and across all analyzed metrics. NB360 reduced plaque density and mean size in brain regions in 5-month but not in 14-month-old mice. Mean plaque-core maturity increased in cortex but decreased in several subcortical structures. LIN5044 was effective in reducing mean plaque size and maturity in many brain areas at 14 months, and on mostly cortical areas at 5 months. Isoctx – isocortex, OLF – olfactory areas, HPF – hippocampal formation, CTX sp – cortical subplate, CNU – caudate nucleus, TH – thalamus, HY – hypothalamus, MB – midbrain, HB – hindbrain, CB – cerebellum

### Local efficacy of therapies with neuroanatomical areas

The effects of the β1 antibody were surprisingly small. In young mice, the increase in plaque density was slightly reduced (gustatory areas and claustrum: p = 0.014) whereas size was marginally increased (claustrum: p = 0.044), and plaque maturity was unaffected. In old mice there was no significant effect (**Figure 2G, S7, S7**). In contrast, NB360 robustly quenched the increase in plaque density and (to a lesser extent) size in 5- month-old mice. The effect on plaque density was most pronounced in subcortical areas (claustrum: p = 0.004) and in ventral and posterior cortical areas including the perirhinal and posterolateral visual area (both p = 0.004), whereas the effect on plaque size was particularly strong in the amygdala and piriform area (both p = 0.024) (**Figure 2F-G, S9**). Plaque maturity (based on the qFTAA/hFTAA fluorescent ratio) was increased in superficial cortical areas (e.g. olfactory areas p = 0.011) but decreased in deep subcortical structures (e.g. amygdala p = 0.017). In old mice, NB360 had no significant effect on plaque density, size and maturity. LIN5044 acted primarily on plaque size, but only marginally on plaque density of old mice (**Figure 2F-G, S10B**). Plaques were smaller in subcortical areas (medial septal complex and amygdala: p = 0.0048 and 0.012 respectively) and cortical areas with a rostro-dorsal emphasis (supplemental somatosensory area: p = 0.0067). The effect of LIN5044 on mean plaque size was more pronounced in old mice, but the spatial distribution of the treatment effect was similar in young and old mice (**Figure 2F-G, S10B**). In contrast, the effect of LIN5044 on plaque density in young mice was less conspicuous.

LIN5044 treatment may influence the fluorescent spectra of plaques and distort maturity analyses. We therefore measured plaque spectra of APP/PS1 mice 3 days after a single injection of LIN5044 or PBS. The emission spectra of plaques were not influenced (**Figure S10A**). In contrast, the cohorts treated for 3 months with LIN5044 showed a massive shift towards increased plaque maturity in old mice (retrosplenial area: p = 0.0014) and to a lesser extent in young mice (supplemental somatosensory area: p = 0.026) (**Figure 2G, S10B**).

### Regional drug – effect analysis based on voxel-level probability distribution

We decomposed atlas-registered brains into spatially registered cubic voxels (15’625 µm^3^) and generated descriptive statistics of plaque density, size and maturity for each voxel. We then assessed the effects of each treatment arm at the single-voxel level (**Figure S5-S6, 3A**). We found that the locales of treatment effectiveness did not coincide with neuroanatomically defined regions. Indeed, assignment by neuroanatomical boundaries failed to capture peaks of regiospecific therapeutic efficacy and overestimated the volume of treatment-affected brain tissue (**Figure 2E**). Remarkably, voxel-level heatmaps of p-values showed that BACE1 inhibition reduced the increase in plaque counts most effectively in the posterior and ventral telencephalon (**Figure 3A**), whereas LIN5044 reduced the increase in the size of plaques primarily in rostro-dorsal areas (**Figure 3A**). These effects were symmetric across the midline and showed sharp boundaries lining the deep cortical layers (LIN5044), the thalamus, and CA3 (NB360). The β1-treated young mice showed patchy reduction in plaque count and size in the brainstem and some decrease in maturity (**Figure S7, S8**). However, these effects were marginal. The effects of β1 in old mice were even less significant (**Figure S7, S8**). Therefore, β1 was excluded from further analyses.

**Figure 3.**
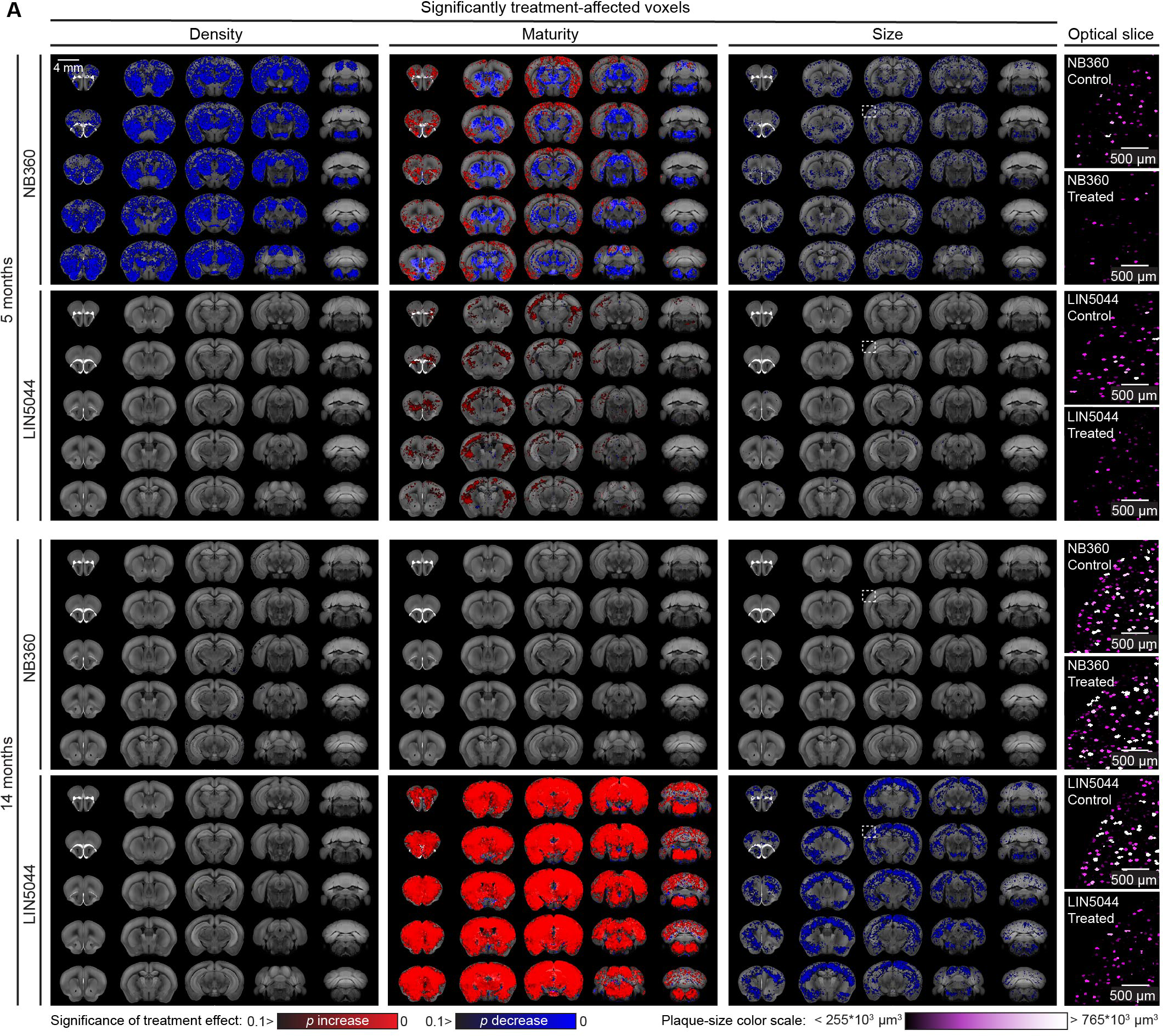
Voxel-based brain analysis reveals temporally distinct anti-Aβ treatment effects across various plaque metrics. (**A**) Each 3-dimensional map of SAVs summarizes all the treated and control samples within a cohort (8-12 samples). These maps reveal regiospecific efficacy unique to each treatment modality. NB360 reduced plaque count and mean size in both cortical and subcortical areas of young (5- month) mice, with little to no effect in older mice (14-month). NB360 also showed profoundly divergent effects on mean plaque maturity between cortical and subcortical regions, which showed increase and decrease, respectively. In older mice, LIN5044 treatment showed a widespread increase in mean plaque maturity, and a significant reduction in mean plaque size across the whole brain. The mean plaque size reduction suggested by the voxel maps was confirmed by looking at segmentations of randomly picked cortical optical slices (white inserts on heatmaps), color-coded by plaque size.

The inferred effects of NB360 and LIN5044 rely on complex computations on terabyte-sized datasets. To intuitively visualize these effects, we randomly selected single cortical mesoSPIM images of atlas-registered brains from each treatment and control groups. Upon segmentation, we color-coded plaques based on their size. **Figure 3A** and **S11** confirm the reduced plaque density and plaque size in NB360 treated young mice and LIN5044 treated old mice, respectively.

### Colocalization analysis reveals little overlap in the regiospecificity of therapies

As a global measure of regiospecific similarity, we counted the overlapping SAVs in all treatment pairs and metrics (plaque density, mean plaque size and maturity) (**Table S4**). Despite a strong colocalization within neuroanatomical boundaries, the LIN5044 and NB360 SAVs appeared to cluster in distinct patterns (**Figure 4A**). The maximal SAV overlap between pairs was <1% or <2.65% (p<0.05 or p<0.1, respectively) indicating that each treatment had a unique voxel-specific fingerprint (**Figure 4A-C, Table S4**). Hypergeometric tests confirmed that the voxel-level overlaps were not significant (p<0.03).

**Figure 4.**
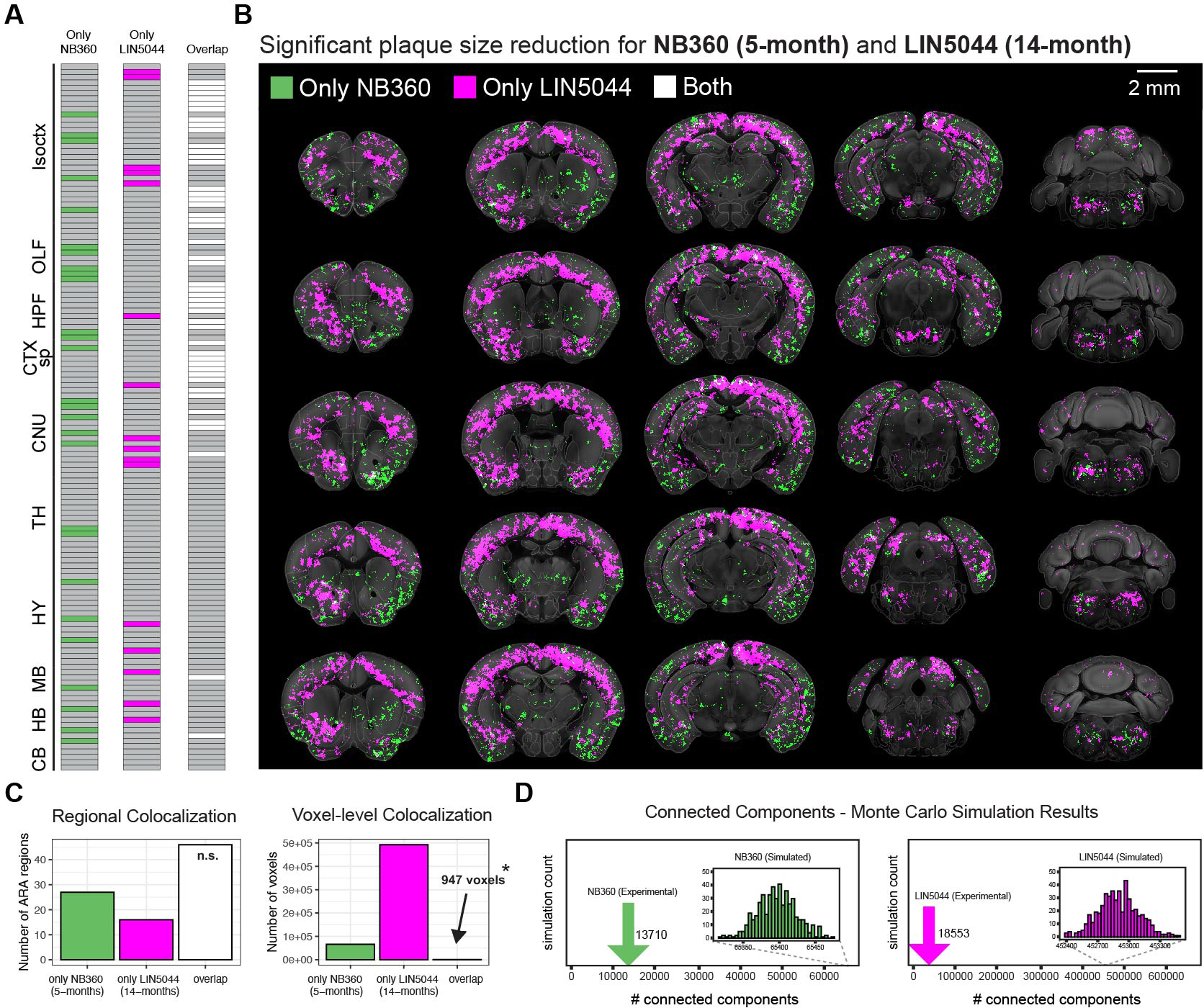
Colocalization of LIN5044 and NB360 voxel-based analysis reveals spatially distinct anti-Aβ treatment effect on plaque size. The distinct significant effect of each of the strongest treatments on plaque size cluster, NB360 5-month and LIN5044 14-month, as well as their overlap, using either (**A**) neuroanatomical boundaries, or (**B**) SAVs. (**C**) The two treatments demonstrate high colocalization based on neuroanatomical boundaries, while the SAVs for plaque-size cluster separately with limited overlap (*p=0.03, hypergeometric test). (**D**) To demonstrate that these individual SAV-clusters are non-random, Monte-Carlo simulations were run which randomly distributed the experimentally-measured number of voxels while the number of connected components was counted. LIN5044 and NB360 SAVs both show at least 5-fold higher clustering than random.

To probe the randomness of such clusters, we measured the number of connected components (neighboring SAVs that are touching each other) in NB360 and LIN5044 treated brains (13’710 and 18’553, respectively). We then generated 500 Monte Carlo simulations with the same number of SAVs than we measured experimentally, but without constraints on their spatial distribution. We found that the number of connected components in the experimental measurements was at least 5-fold lower than in the simulations, indicating that the SAVs are grouped into distinct, spatially confined clusters (**Figure 4D**).

### Regiospecificity is not due to pharmacokinetics or beta-secretase activity

To test if the regiospecificity was caused by differential penetration of therapeutic compounds, we determined the biodistribution of β1, NB360 and LIN5044 by dissecting brains into 8 standard regions (**Figure S12A**). NB360 levels were measured 1 hour after oral administration (Neumann, Rueeger et al. 2015). Since antibodies have long half-lives (Vieira and Rajewsky 1988) and limited blood-brain barrier penetration, brain levels were measured 6 and 24 hours after intraperitoneal injection. As the pharmacokinetic properties of LIN5044 are unknown, we measured brain levels 2 and 6 hours after intraperitoneal administration. There was no difference in regional NB360 levels. LIN5044-treated brains showed higher levels in the brainstem and cerebellum (p<0.034 and 0.024, respectively) after 2 hours, but its distribution became homogeneous after 6 hours (**Figure S12B-D**). β1 showed higher levels in the brainstem after 6, and in the brainstem and the cerebellum after 24 hours (**Figure S12E, F**). However, there was no difference in antibody levels between diencephalic and telencephalic regions. Pooled non-specific recombinant IgG was used for control and did not accumulate in any brain region at 24 hours (**Figure S12G**). Hence regional pharmacokinetic differences do not explain the region-specific drug effects. We also tested the abundance of Aβ and BACE1 in 8 brain regions (**Figure S12A**) biochemically (**Figure S13**). We could not differentiate between brain regions biochemically; however, both LIN5044 and NB360 reduced the amount of Aβ monomers compared to control mice (**Figure S13**).

### Genetic markers revealed by aligning the Q3D output to gene-expression atlases of the brain

The findings above suggested that the local heterogeneity of drug efficacy may be controlled by intrinsic properties of the host brain. We therefore compared the plaque-size SAVs of NB360 and LIN5044 to a gene-expression atlas reporting whole-genome expression at >30’000 spots of the mouse brain(Ortiz, Navarro et al. 2020) (**Figure 5A**). We calculated the mutual information (MI), a similarity metric describing the non-linear interdependence of random variables. We also created an online application for browsing this data (https://fgcz-shiny.uzh.ch/SPAGEDI/). As expected, most genes showed low MI scores with either treatment (**Figure 5B**). The MI of *Thy1*, whose promoter was used to drive APP/PS1 expression, ranked at the 99^th^ percentile of 23371 genes for both LIN5044 and NB360 (MI: 8.06×10^-4^; NB360 5.32×10^-4^, respectively). Similarly, *Bace1* expression correlated highly with the efficacy of its inhibitor NB360 (MI: 2.22×10^-4^, 96^th^ percentile), but not with LIN5044 (1.37×10^-4^, 81^st^ percentile) (**Figure 5C**). These results confirm that the morphological-genetic analyses presented here identify sensitively and reliably the genetic networks controlling the efficacy of amyloid removal therapies.

**Figure 5.**
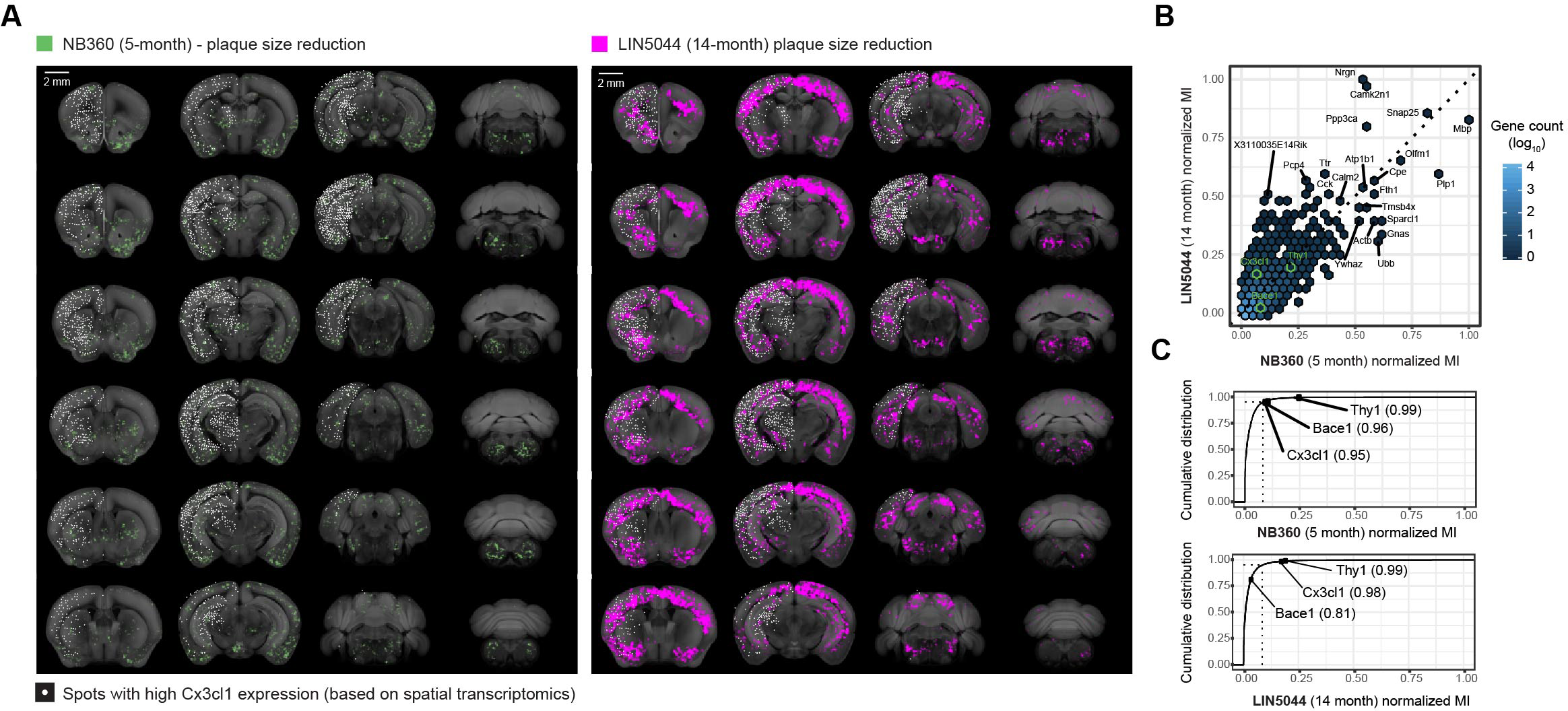
Genetic markers may predict local drug responsiveness. (**A**) SAV plaque-size-reduction maps for NB360-5-month and LIN5044-14-month effects overlaid with data from a spatial transcriptomics database, with the Cx3cl1 gene shown as an example. Spots with high Cx3cl1 expression (white) show more overlap with the LIN5044 effect than with NB360. Calculating the mutual information between each SAV and gene expression pair allows for the discovery of candidate genetic markers that predict each drug’s responsiveness. (**B**) Normalized Mutual Information (MI) of genes above the 95^th^ percentile (ranked by MI, [n=1169]) for either NB360 or LIN5044. (**C**) MI between SAV and gene expression ranked cumulatively by MI (23371 genes). Cx3cl1, a neuron-borne microglia chemoattractant, is a higher ranked genetic marker for LIN5044 responsiveness. Dashed line: genes above the 95^th^ percentile of MI.

The effect of LIN5044 showed a high MI (7.23×10^-4^, 98^th^ percentile) with Cx3cl1, a microglial chemoattractant mostly expressed by neurons (Tarozzo, Bortolazzi et al. 2003) (NB360, Cx3cl1 MI: 1.95×10^-4^, 95^th^ percentile) (**Figure 5**), potentially suggesting that the effects of LIN5044 are stronger in areas of high microglia recruitment. To test this, we quantified Iba1^+^ microglia in cortical areas with strong or weak LIN5044 effects. Regions strongly affected by LIN5044 showed significantly higher microglia counts (**Figure S14**).

## Discussion

Brain-wide analysis of Aβ plaque load, as shown previously(Liebmann, Renier et al. 2016), requires reliable brain clearing and amyloid staining (Richardson, Guan et al. 2021). However, current hydrogel-based methods for whole-brain clearing are slow and often inhomogeneous. We have solved these limitations by insulating the anodic from the cathodic detergent reservoir, thereby constraining the lipid-clearing ion flow through the sample. This design enabled whole-brain clearing within 14 hours (Chung, Wallace et al. 2013, Tomer, Ye et al. 2014, Richardson and Lichtman 2015), and electrophoresis of qFTAA/hFTAA resulted in homogenous plaque staining within 2 hours. Comparisons between the Q3D pipeline, including light-sheet acquisition (3.26 × 3.26 × 6.5 µm resolution), with conventional staining of histological sections showed that Q3D reliably visualizes Aβ plaques within mouse brains. The precision of Q3D resulted in highly consistent plaque counts among age-matched APP/PS1 mice (standard deviation: 2.5-3.3%).

The β1 antibody showed only marginal effects in young APP/PS1 mice and no effect in old mice, consistent with previous reports (Pfeifer, Boncristiano et al. 2002, Balakrishnan, Rijal Upadhaya et al. 2015). The marginal effect was primarily visible in the brainstem, where higher β1 antibody drug concentrations were measured (**Figure S11**). It was shown that β1 injections result in target engagement (Balakrishnan, Rijal Upadhaya et al. 2015), thus the limited effect of the β1 antibody may be explained by its low bioavailability relative to the high abundance of Aβ. In contrast, both LIN5044 and NB360 reduced amyloid load more effectively. This provided the first evidence that polythiophenes can be efficacious against Aβ. However, we unexpectedly found that the activities of LIN5044 and NB360 were highly divergent in many ways. As our colocalization analysis revealed, the spatial efficacies of LIN5044 and NB360 were almost mutually exclusive, being more effective against either rostro-dorsal or ventrocaudal amyloid deposits, respectively. Moreover, LIN5044 was more effective in reducing the growth of existing plaques, rather than their total number.

LIN5044 was also more effective in aged mice than in younger mice. In contrast, BACE1 inhibition was most effective in reducing plaque numbers in young mice, yet affected their growth to a lesser extent. The reduction in plaque burden showed steep ventrocaudal (NB360) and dorsorostral (LIN5044) gradients whose boundaries did not correspond to defined neuroanatomical areas or vascular territories. Thus, the voxel-level analysis was crucial in finding clusters of treatment significance beyond conventional anatomical boundaries.

The strong Aβ-monomer reducing effect of BACE1 decreases plaque loads in young, but not in old, mice, suggesting that distinct amyloid species are contributing to plaques at different ages. While primary nucleation is mainly dependent on Aβ concentration (Hellstrand, Boland et al. 2010, Burgold, Filser et al. 2014) and is highly sensitive to BACE1 inhibition (Brendel, Jaworska et al. 2018, Peters, Salihoglu et al. 2018), secondary nucleation (Cohen, Linse et al. 2013) may become more important in aged mice. In prion diseases, polythiophenes (including LIN5044) slow disease progression by reducing the number of seeds for secondary nucleation. In prion diseases, polythiophenes (including LIN5044) slow disease progression by reducing the number of seeds for secondary nucleation. This is achieved by polythiophenes binding and stabilizing amyloid fibrils, resulting in reduced fibril fragmentation as a source of seeds (Margalith, Suter et al. 2012, Herrmann, Schutz et al. 2015) . Similarly, Aβ fibril hyper-stabilization could explain the effect of LIN5044 in APP/PS1 mice. The remarkable efficacy of LIN5044 in aged mice. may relate to the dependency of secondary nucleation onto the concentration of seeds, which increases with age. Accordingly, LIN5044 in older mice primarily reduced mean plaque sizes (**Figure 2G**), all while increasing the number of smaller plaques (**Figure 2F**), providing further evidence that LIN5044 reduces additive plaque growth. Potentially, smaller plaques might be a consequence of an increase in plaque compaction upon LIN5044. The stronger effect of LIN5044 in old mice suggests that it preferentially targets tightly packed plaques, which become more abundant with time. Pharmacokinetic analyses did not support the notion that differential bioavailability may account for the regiospecific activities of NB360 and LIN5044 (**Figure S11**). Also, we did not detect regional differences in endogenous Aβ and BACE1 levels. Because of its inherent fluorescence, LIN5044 might also influence the spectral properties of plaques. We therefore compared plaque spectra in a mouse injected i.p. with LIN5044 (0.4 mg) and one injected with PBS. No differences were detected, suggesting that the effect of LIN5044 on plaque maturity is not artefactual (**Supplementary Fig. 9**).

By aligning SAV heatmaps to a spatial transcriptomic atlas (Ortiz, Navarro et al. 2020) we identified a strong spatial similarity between *Bace1* expression and the regiospecific efficacy of NB360, but not of LIN5044. *Thy1*, the promoter driving the APP/PS1 mouse model, ranked highly for both treatments, providing validation for the unsupervised approach chosen here. LIN5044 efficacy showed high spatial similarity with transcription maps of CX3CL1, a neuron-borne microglial chemoattractant (Szepesi, Manouchehrian et al. 2018). The regional variation of CX3CL1 expression may result in differential recruitment of microglia, thereby contributing to the clearance of LIN5044-intercalated amyloid. Indeed, follow-up experiments confirmed that the areas where LIN5044 had a stronger effect also harbored more microglia in APP/PS1 mice. The cognate microglial receptor CX3CR1 is highly expressed at late AD stages (Chen, Lu et al. 2020), which may partly explain the potency of LIN5044 in old mice. As shown here, correlating spatial transcriptomic atlases to therapy atlases can be used as a tool to generate hypotheses for treating Aβ pathology. However, when using such an approach one needs to take into consideration that gene expression and protein localization might not coincide. In the future, such spatial analyses could benefit from incorporating proteomic data.

Our results suggest that the regional variation and age-dependence of anti-amyloid drug efficacy could considerably influence clinical trial efficacy. Since LIN5044 and NB360 affect mostly non-overlapping areas of the brain, combinatorial regimens may synergistically protect larger brain volumes from amyloid deposition. Notably, the computational methods developed for Aβ amyloid quantification can enhance the statistical power of the in vivo assessment of anti-Aβ drugs and can be adapted to a broad range of protein aggregation diseases.

## Materials and Methods

### Animal treatments and tissue preparation

All animal experiments were carried out in strict accordance with the Rules and Regulations for the Protection of Animal Rights (Tierschutzgesetz and Tierschutzverordnung) of the Swiss Bundesamt für Lebensmittelsicherheit und Veterinärwesen and were preemptively approved by the Animal Welfare Committee of the Canton of Zürich (permit 040/2015). APP/PS1 male and female mice were either treated with NB-360 (Neumann, Rueeger et al. 2015) BACE1-inhibitor (Novartis) orally (0.5 g inhibitor/kg chow, ∼6 g chow/day/mouse = 3mg inhibitor/day/mouse), with the β1 monoclonal IgG2a antibody recognizing the human specific EFRH tetrapeptide of amino acids 3 - 6 of Aβ (Paganetti, Lis et al. 1996, Pfeifer, Boncristiano et al. 2002, Balakrishnan, Rijal Upadhaya et al. 2015) (Novartis) (0.5 mg/week, once/week in 200µl phosphate buffered saline (PBS) intraperitoneally), or with the amyloid intercalator LIN5044 (0.4 mg/week, once/week in 100 µl PBS) based on previous reports. Because the pharmacokinetics of LIN5044 was unknown, the dose was selected based on previous work (Herrmann, Schutz et al. 2015). Control mice were treated with control chow, pooled recombinant non-specific IgG, and PBS, respectively. The ages of NB-360, β1 antibody, and LIN5044 treated mice were 353 ± 21, 313 ± 9, and 308 ± 7 days, respectively (groups of old mice), as well as 61 ± 3 days, 59 ± 2 and 65 ± 2 days, respectively (groups of young mice) (Table S1). After treatments were completed, mice were deeply anaesthetized with ketamine and xylazine, then transcardially perfused with ice-cold phosphate-buffered saline (PBS) followed by a hydrogel monomer mixture of 4% acrylamide, 0.05% bisacrylamide and 1% paraformaldehyde(Yang, Treweek et al. 2014). Brains were harvested and further incubated passively in the hydrogel mixture for 24 hours. The hydrogel was degassed and purged with nitrogen, followed by polymerization at 37 °C for 2.5 hours. Samples were either stored in PBS or clearing solution until Q3D clearing.

For generating the whole-brain vascular images (**Figure S3A**), Claudin5-GFP (Gensat.org. Tg(*Cldn5*-GFP)) mice were processed as described above.

### Tissue clearing

Brains were cleared with focused electrophoretic tissue clearing (FEC) in a custom-built chamber in 8% clearing solution (8% w/w sodium dodecyl sulphate in 200 mM boric acid, pH 8.5). Standard settings were 130 mA current-clamped at a voltage limit of 60V, at 39.5 °C. Clearing time varied between 6-14 hours for a whole mouse brain. Tissue clarity was determined by visual inspection. The polarity of the electrodes was switched after approximately 50% of the total clearing time. Transparency was assessed by visual inspection. The clearing solution was circulated from a buffer reservoir of 250 ml. After clearing a brain, 60 ml of clearing buffer was exchanged with fresh buffer before starting to clear the next sample. Q3D-clearing chambers were 3D-printed, and clearing was done in an incubator at 39.5 °C. For details on the chamber design, see supplementary material and model repository (**Figure S1**) (Kirschenbaum 2020).

### Relative electrical resistivity measurement between brain tissue and buffer

For electrical resistivity measurements comparisons, mouse brains were first fixed in 4% paraformaldehyde, then measured in PBS. Platinum electrodes (1.5 × 0.2 mm) were mounted at a distance of 2 mm, and a constant voltage of 30 V was applied. Current measurements were used to calculate the resistance between the electrodes. Brain tissue resistance was measured by sticking the electrode-pair into the brain at 3 different locations (frontal cortex, occipital cortex and brainstem). At each location, current values were measured three times. The resistivity in the buffer was measured nine times. These measurements resulted in a ∼1200-Ohm resistance in the brain and ∼300-Ohm resistance in the buffer. As the electrode setup and the voltages were constant for every measurement, the relative resistivity was calculated as the ratio of the two resistivity measurements, resulting in a resistivity ratio of 1:4 (buffer:brain).

### Comparison between Q3D and CLARITY

Hydrogel-embedded brains from 3-month-old mice were either passively cleared with 8% clearing solution for 24 hours at 39°C (n = 3) or for 4-6 hours with either Q3D or CLARITY (Chung, Wallace et al. 2013) clamped at 130 mA at 60V at 39 °C (four active clearing groups, each n = 3). After clearing, brains were washed with PBS, followed by refractive index-matching and mounting in quartz cuvettes. Samples were illuminated with a ∼4-5 µm wide 647-nm laser beam, calibrated to 0.35-mW power. Each brain was illuminated stereotypically from the dorsal end at 10 aligned points with 0.5 mm spacing along the rostro-caudal axis, with the same incumbent laser power (as measured through the imaging medium and cuvette). This was done in 3 parallel lines (one in the midline, and a line 2 mm left and right from it), resulting in 3×10 measurement points per brain (**Figure S1F**). The point pattern was defined by starting from the lambdoid fossa. The transmitted light was measured at each point with a digital optical power meter (Thorlabs, PM100D, Compact Power and Energy Meter Console, Digital 4″ LCD; Thorlabs S130C photodiode sensor). The mean and standard error of the mean for all points for each sample was plotted (**Figure S1F-G**). We also plotted the mean transmitted light on every rostro-caudal level, resulting in 10 datapoints summarizing the 3 parallel lines (two hemispheres and the midline) (0.35mW input power, 647nm wavelength, 3 brains/group, 30 measurement points/brain, **Figure 1C and S1**). These 10 points per sample were submitted to a one-way ANOVA (α = 0.05), which was corrected with Tukey’s test for multiple comparisons.

### Electrophoretic staining with Q3D

Histochemistry by iontophoretic tissue staining was initially done by layering paraffin and agarose around the sample in order to limit current flow to the tissue. Later, Q3D staining chambers were 3D-printed (**Figure S2A-B**). 50 mM tris and 50 mM tricine at a pH of 8.5 was used as the electrophoresis buffer. Tests in native polyacrylamide gel electrophoresis were followed by silver staining (Thermo Scientific Pierce Silver Stain Kit, #24612) that showed, as expected, that electrophoretic mobility is highly dependent on the pH and buffer (**Figure S2D-G**). Amyloid plaques were stained with a combination of luminescent conjugated polythiophenes (LCP), heptamer-formyl thiophene acetic acid (hFTAA), and quadro-formyl thiophene acetic acid (qFTAA). The combination of these dyes was used for the discrimination of neuritic plaques (Nystrom, Psonka-Antonczyk et al. 2013) at different maturation states(Rasmussen, Mahler et al. 2017).

### Refractive index matching

Brains that were cleared and stained with Q3D were refractive index (RI) -matched to 1.46 with a modified version of the refractive index matching solution (Yang, Treweek et al. 2014) by including triethanolamine (tRIMS). tRIMS was made by mixing Histodenz (Sigma #D2158) (100 mg), phosphate buffered saline (75 ml), sodium azide (10% w/v, 500 μl), tween-20 (75 μl) and triethanolamine (42 ml). tRIMS maintained the RI while reducing the amount of Histodenz required and improving transparency. After prolonged air and light exposure, tRIMS tends to undergo browning; however, in air-tight tubes at 4 °C samples remain stable for at least two years.

### Antibody and polythiophene staining

Slices from formalin fixed and paraffin embedded brain tissue from a 13-month-old APP/PS1 mouse were stained for Aβ plaques. Slices were stained with mouse anti-human Aβ_17-24_ antibody (4G8, Biolegend SIG-39220) after antigen retrieval with 10% formic acid. Slices were blocked with M.O.M. Kit (BMK-2202) and the primary antibody was detected with Alexa-594 conjugated goat anti-mouse IgG (Invitrogen A-11005, 1:1000 dilution). Alternatively, slices were stained with Aβ N3pE rabbit anti-human antibody (IBL 1A-018, 1:50 dilution) after 10% formic acid antigen retrieval, followed by blocking with 10% goat serum, detected with Alexa-594 conjugated goat anti-rabbit IgG (Invitrogen A-11037, 1:500 dilution). Both antibody stainings were followed by staining with qFTAA (0.75 μM in PBS) and hFTAA (3 μM in PBS) for 30 minutes, followed by diamidino-phenylindole (DAPI) staining. Slices were imaged with a Leica SP5 confocal microscope with a 10x air objective (numerical aperture 0.4). The qFTAA and hFTAA stainings were imaged by exciting both at 488 nm and collecting emission between 493 - 510 nm and 530 – 579 nm, respectively. The dynamic range of images was adjusted consistently across stainings and images (3.57 µm/pixel) were median filtered with ImageJ (pixel radius 0.5).

To test the effect of the LIN5044 treatment on the qFTAA+hFTAA plaque maturity analysis, 12-month-old APP/PS1 mice (n=2) were injected i.p. with LIN5044 (0.4 mg in 100µL PBS) (n=1) and PBS (100 µL) (n=1). After 3 days mice were deeply anaesthetized with ketamine and xylazine and transcardially perfused first with ice cold PBS, followed by 4% paraformaldehyde. Brains were harvested and were further incubated in the paraformaldehyde solution for 24 hours. Brains were then incubated in 30% sucrose in PBS for two days at 4 °C. Next, brains were snap-frozen and stored at -80 °C overnight. Brains were embedded in tissue freezing medium (Leica biosystems), and 20 μm coronal slices were cut with a cryostat and mounted on Super-frost micro-slides. Tissue slices were fixed in 10% formalin overnight and rehydrated by dipping them in consecutive baths of 99% ethanol, 70% ethanol, dH2O, and PBS, 10 min in each. The tissue sections were allowed to dry under ambient conditions. For polythiophene staining 200 µL of hFTAA and qFTAA solution (20 µM) were added onto the tissue sections to cover them and slices were incubated for 30 min at room temperature. Sections were rinsed in a PBS bath for 10 min, followed by nuclear staining with DAPI. Tissue sections were dried under ambient conditions, followed by mounting with fluorescence mounting medium (DAKO). Slides were imaged with a Leica SP5 Confocal microscope. Nuclei and plaques were imaged with a 10X/0.25 NA dry objective, using the following settings: 405/30 nm excitation/emission filters for DAPI (nuclei) and 498-520nm (compact amyloid) and 565-605nm (looser amyloid) for LCPs. Laser power was set on 10% for all the conditions and a line average of 96 was used for acquisition. Three 8-bit images were recorded from each sample. Ten plaques per image were measured for their fluorescent intensity by quantifying the maximum intensity of a line drawn through each individual plaque with Fiji. The mean intensity of ten plaques per image was compared with a 2-tailed T-test between the two conditions (LIN5044 or PBS treated). For microglia immunohistochemistry, fixed brains of LIN5044 treated transgene-negative littermates from APP/PS1 nests (n = 3) were embedded in paraffin. 4-μm-thick paraffin sections (3 sections per mouse) were deparaffinized through a decreasing alcohol series. Slices were stained with Iba-1 antibody (1:1000; Wako Chemicals GmbH, Germany) and detected using an IVIEW DAB Detection Kit (Ventana). Sections were imaged using a Zeiss Axiophot light microscope. For the quantification of the Iba-1 staining, in every slice two regions of interest were selected in the cortex. The four regions of interest were selected representing 2-2 cortical areas with either high or no LIN5044 therapeutic effect (**Figure 3A, 5D-E**). Pixels in the regions of interest were classified and counted as microglia (Iba-1 positive) or background (Iba-1 negative) with a manually trained (trained on three images) pixel classifier in ILASTIK (https://www.ilastik.org/), and ImageJ. Hypothesis testing was done with a 2-tailed T-test.

### Drug distribution measurements

NB360 was administered orally in three male C57BL/6 black mice at 5mg/kg body-weight dose-level at 0.5 mg/ml in water with 0.5% methylcellulose and 0.1% Tween-80. Based on preceding pharmacokinetic studies (Neumann, Rueeger et al. 2015), brains were harvested one hour later, followed by homogenization in water and acetonitrile precipitation. NB360 levels were measured with tandem mass spectrometry with electrospray ionisation. LIN5044 was administered intraperitoneally into male C57BL/6 black mice at 16 mg/kg body-weight dose-level at 4 mg/ml in PBS. As preceding pharmacokinetic studies were not available, two- and six-hour incubation timepoints were chosen, each with three mice. Brains were homogenized in water and precipitated with methanol. LIN5044 levels were measured with high pressure liquid chromatography - tandem fluorescence detection. Brain regions’ drug levels were compared with one-way ANOVA (α = 0.05) for each compound and timepoint separately (NB360, LIN5044 2 hour, LIN5044 6 hour); multiple comparisons were corrected for with Tukey’s test.

### Antibody brain-distribution measurements

C57BL/6 mice were injected one-time with either β1 antibody (n=6), as a control with pooled recombinant non-specific IgG (n=2) (0.5 mg in 200 µl intraperitoneally), or with no injection (n=1). β1 injected mice were sacrificed after 6 (n=3) or 24 hours (n=3), while control mice were sacrificed 24 hours after injection. Amyloid β Protein Fragment 1-42 (A9810, Sigma) was diluted at 1 µg/mL in PBS and passively absorbed on multiwell plates (SpectraPlate-384 HB, Perkin Elmer) overnight at 4 °C. Plates were washed three times in 0.1% PBS-Tween 20 (PBS-T) and blocked with 80 μl per well of 5% skim milk (Migros) in 0.1% PBS-T, for 2 h at room temperature. β1-antibody and pooled recombinant IgG were used as positive and negative controls, respectively. Blocking buffer was discarded, and both samples and controls were dissolved in 1% skim milk in 0.1% PBS-T for 1 h at 37°C. 2-fold dilutions of β1-antibody, starting at a dilution of 1000 ng/ml in 1% skim milk and in 0.1% PBS-T were used for a calibration curve. Goat polyclonal anti-mouse antibody (1:1’000, 115-035-062, Jackson ImmunoResearch) was used to detect murine antibodies. Chromogenic reaction was induced by addition of TMB Stabilized Chromogen (SB02, Thermo Fisher Scientific) and stopped by addition of 0.5 M H_2_SO_4_. Absorbance was read at *λ* = 450 nm. Unknown β1-antibody concentrations were interpolated from the linear range of the calibration curve using linear regression (GraphPad Prism, GraphPad Software).

### Western Blots of Aβ and BACE1

Brain hemispheres from mice treated with LIN5044 (0.4 mg/week, once/week in 100 µl PBS) (n=1) or PBS (n=1) intraperitoneally, or with NB360 chow (0.5 g inhibitor/kg chow, ∼6 g chow/day/mouse = 3mg inhibitor/day/mouse) (n=1) or control chow (n=1) for 3 months, were dissected into 8 anatomical regions: rostro dorsal, rostro ventral, medio dorsal, medio ventral, caudo dorsal, caudo ventral, brain stem, cerebellum (**Figure S19A**). Each region was homogenized using Ribolyser for 5 min in 500uL lysis buffer (140mM NaCl, 20mM TrisHCl, pH 7.5, protease inhibitors [complete Mini, Roche], phosphatase inhibitors [PhosphoSTOP, Roche] in PBS), and centrifuged at 15000g for 30 min at 4°C. Subsequently, the supernatant of each sample was isolated and the pellet was resuspended in 200 uL of lysis buffer with the addition of 0.5% SDS to obtain the insoluble fraction. 2uL of Dithiothreitol (DTT) was added to 20uL of each fraction. Samples were loaded on a SDS-PAGE (Novex NuPAGE 4-12% Bis-Tris Gels). After electrophoresis, gel was transferred to iBlot I (Invitrogen) and transferred onto Polyvinylidene difluoride (PVDF) membrane. Membranes were blocked in 5% Sureblock for 1 h at room temperature followed by incubation at 4 °C overnight with 1:1000 dilution of the following primary antibodies: mouse monoclonal to human amyloid beta 1-16, clone 6E10 (Sigma) or rabbit polyclonal to BACE1 (abcam ab2077). Membranes were washed 3x (10 min each) with PBS-Tween (0.1%) followed by incubation with HRP-tagged secondary antibody (Peroxidase-Goat Anti-Mouse IgG (H+L) (#62-6520) or Peroxidase-Goat Anti-Rabbit IgG (H+L) (#111.035.045); 1h at room temperature) and further washes (3x, 10 min). Membranes were developed with Luminata Crescendo (Millipore) and images were acquired using Fusion Solo S (Vilber).

### Beta-secretase activity assay

Homogenates from the 8 anatomical regions (see Western Blots in the method section) were used to assess the difference in beta-secretase activity in each anatomical region using the Beta-Secretase Activity Fluorometric Assay Kit (Merck, MAK237).

### Whole-brain imaging

Whole brain images were recorded with a custom-made selective plane illumination microscope (mesoSPIM)(Voigt, Kirschenbaum et al. 2019). SPIM imaging was done after clearing and refractive index matching. The laser/filter combinations for mesoSPIM imaging were as follows: for qFTAA at 488 nm excitation, a 498 - 520 nm bandpass filter (BrightLine 509/22 HC, Semrock / AHF) was used as the emission filter; for hFTAA at 488 nm excitation, a 565 - 605 nm bandpass filter (585/40 BrightLine HC, Semrock / AHF) was used. Transparent whole-brains were imaged at a voxel size of 3.26 × 3.26 × 3 µm^3^ (X × Y × Z). For scanning a whole brain, 16 tiles per channel were imaged (8 tiles per brain hemisphere). After the acquisition of one hemisphere, the sample was rotated and the other hemisphere was then acquired. The entire process resulted in typical acquisition times of 2-3 hours, followed by stitching(Bria and Iannello 2012). Data accumulated from one brain ranged around 600 GB in size. Further technical details of the mesoSPIM have been previously reported(Voigt, Kirschenbaum et al. 2019).

### Computational and statistical analysis

The following computations were performed using custom scripts written in Python and R(Dadgar-Kiani 2020) as well as existing third-party libraries (Table 2). The 2-channel (498–520 nm and 565-605 nm) substacks for each brain hemisphere were first stitched together with Terastitcher (Bria and Iannello 2012). The result was downsampled from the acquired resolution (3.26 µm lateral, 3 µm depth) to an isotropic 25 µm resolution and then registered to the Allen Institute 25 µm average anatomical template atlas (Wang, Ding et al. 2020). This was performed automatically using a combination of affine and b-spline transformation registrations with a mutual information similarity metric, using parameters influenced from a previous study performing mouse whole-brain fluorescence quantification (Renier, Adams et al. 2016). The resulting pairs of transformations were used in subsequent steps to transform coordinates in the raw data space to the template atlas space (**Figure S4D**).

The 565-605 nm channel at its original resolution was used to determine the locations of aggregates of amyloid-β stained with qFTAA and hFTAA. A random forest classifier was used to classify each voxel as either “belonging to a plaque” or “background”. This classifier was generated using the open-source Ilastik framework (Berg, Kutra et al. 2019), and trained on a random subset of data (random stacks, 3 stacks [187×176×1242 pixels] picked from every experimental group) that was separately annotated by two neuropathologists. Amyloid-β aggregates were considered to be the individually connected components from this binarized volume. The three-dimensional center of mass and total volume was then calculated for each component. Connected components with a volume below a global threshold were considered noise and ignored. The centers of mass were used to look up the peak fluorescent intensity of each plaque in the 498-520 nm (qFTAA) channel. Plaque maturity was calculated for each plaque as its peak intensity in the 498-520 nm (qFTAA) channel divided by its peak intensity in the 565-605 nm (hFTAA) channel (Nystrom, Psonka-Antonczyk et al. 2013) (**Figure 2C**).

After downsampling each aggregate center to 25-µm resolution and applying the optimized registration transformation, the number of aggregates were counted at each voxel in this atlas space (**Figure S4E**). Smoothed heatmaps were generated by placing a spherical ROI with 15-voxel diameter (= 375µm) at each voxel and summing the plaque counts within the ROI. This ROI diameter was set to match the mean spatial jacobian-matrix determinant of the previously-registered b-spline transformation across all samples. This method for smoothing and accounting for variable registration quality has also been described in a previous whole-brain study (Renier, Adams et al. 2016). Voxel-level statistics across treated and control brains involved running a two-sided t-test at each heatmap voxel across the two groups. Each voxel p-value was adjusted using the Benjamini-Hochberg method (Benjamini, Drai et al. 2001). These adjusted p-value maps were then binarized with a threshold of 0.05 or 0.10 for subsequent analysis or visualization.

The transformed locations of each plaque were also further grouped into 134 different anatomically segmented regions in the Allen Reference Atlas (Wang, Ding et al. 2020) for further statistical analysis between longitudinal groups (**Figure S4F, S15**). Similar heatmap generation, voxel statistics, and regional statistics were performed for two other metrics: the mean plaque volume and the mean plaque maturity.

The voxel-level statistical map for each treatment group was also compared against a spatially-resolved transcriptomics database consisting of 23371 genes across 34103 locations throughout the brain (Ortiz, Navarro et al. 2020). Since the data from this study and the spatial transcriptomic database were both in the ARA coordinate space, it was possible to spatially compare each gene’s expression map with each treatment group’s binarized p-value map using a similarity metric. In this case, mutual information was used as the similarity metric since it will detect any sort of linear or non-linear relationship between these two discrete datasets. In order to rank the relevancy of genes in contributing to the spatial map of plaque removal efficacy for a particular compound, we generated a ranked list of genes for each treatment group, sorted by their corresponding mutual information score. We also created an online browser for the MI database (https://fgcz-shiny.uzh.ch/SPAGEDI/)(Dadgar-Kiani 2021).

*Key Resources are summarized in supplemental table 2* (**Table S5**)*. See also References(Bradski 2000, Klein, Staring et al. 2010, Millman and Aivazis 2011, Furth, Vaissiere et al. 2018)*

## Acknowledgements

We thank Dr. Giulia Miracca, Dr. Todd E Golde and Dr. Soyon Hong for critical reading of the manuscript, Dr. Ulf Neumann for providing NB360 and β1, Dr. Michael B. Smith for help with designing 3D-printed parts, and Dr. Asvin Lakkaraju for help with biochemical measurements. Scientific sketches were used from SciDraw.io doi.org/10.5281/zenodo.3925971, doi.org/10.5281/zenodo.3925911, doi.org/10.5281/zenodo.3926119.

## Authors contributions

D.K., J.H.L., and A.A. designed the study. D.K. conducted the experiments, acquired and analyzed the data. D.K. and A.A. prepared the figures and wrote the paper. F.F.V. designed and built the imaging platform. E.D.K. developed the computational analysis pipeline, analyzed data, wrote the paper and created figures. F.C. conducted animal experiments and imaging. O.B. worked on tissue clearing. P.N. and H.S. developed polythiophenes. K.J.F. measured antibody concentrations with ELISA. P.P. developed anti-amyloid β antibodies. F.H. helped develop the imaging platform. J.H.L. and A.A. supervised personnel and wrote the paper.

## Data availability

All data needed to evaluate the conclusions in the paper are present in the paper and/or the Supplementary Materials. The computational code developed for this paper is available upon request. The imaging data is available upon request.

## Figure Legends

**Supplementary Figure 1.**
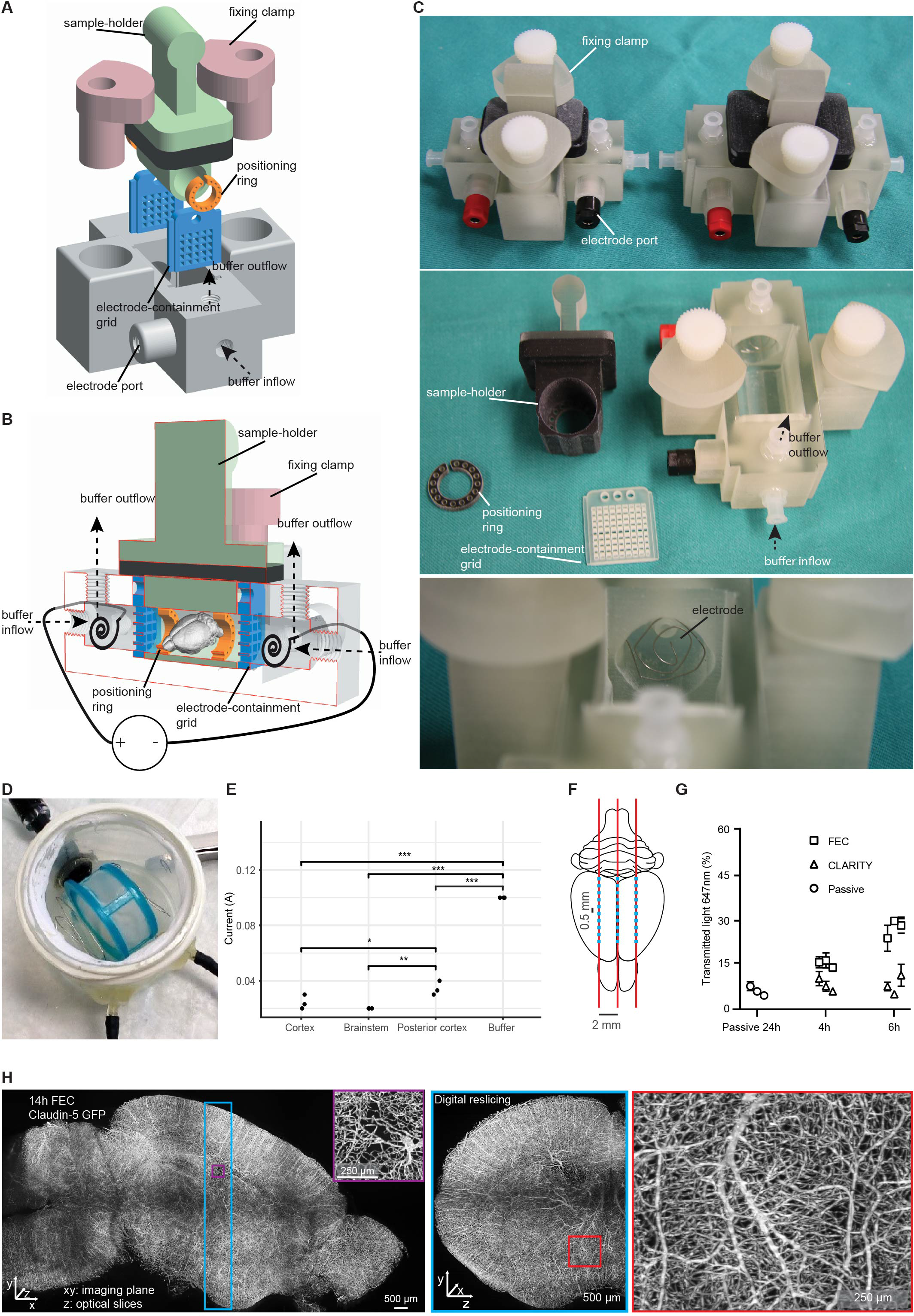
Focused electrophoretic tissue clearing (FEC) chambers allow for improved tissue clearing efficiency. (**A-C**) Computer designed and 3D-printed Q3D-clearing chambers. (A) Rendering and (B) cross-section of a chamber. The clearing solution is circulated using two independent, electrically isolated circuits around each electrode. The tissue is contained in a sample-holding plunger (green), and is separated from the electrodes by small nets (blue). The plunger is sealed by a rubber ring (black). Samples are positioned within the tube with rings (orange). (**C**) 3D printed FEC clearing chambers in different sizes. (**D**) Reproduction of a CLARITY chamber for comparison. (**E**) Electric current measurements in different parts of 4% paraformaldehyde fixed brain in PBS and in PBS buffer (*p<0.05, **p<0.01, ***p<0.001). (**F-G**) CLARITY and FEC were compared by measuring the light transmitted through the brain in 30 points (f, blue dots). Measurement points were defined along 3 parallel lines 2 mm apart, one in the midline (red lines). Then, for every measurement the laser beam was moved in 0.5 mm steps rostrally starting from the lambdoid fossa. (**G**) Higher transparency in FEC-cleared samples after 4 and 6 hours. Each datapoint represents the mean of all 30 measurements in one mouse brain (n=3 per group). (**H**) The brain vasculature of Claudin5-GFP mice after 14 hours of clearing was imaged sagittally by mesoSPIM. Digital reslicing and visualization by maximum-intensity projection demonstrates that focused clearing resulted in highly uniform signals. Darkened areas are due to stitching/vignetting artifacts.

**Supplementary Figure 2.**
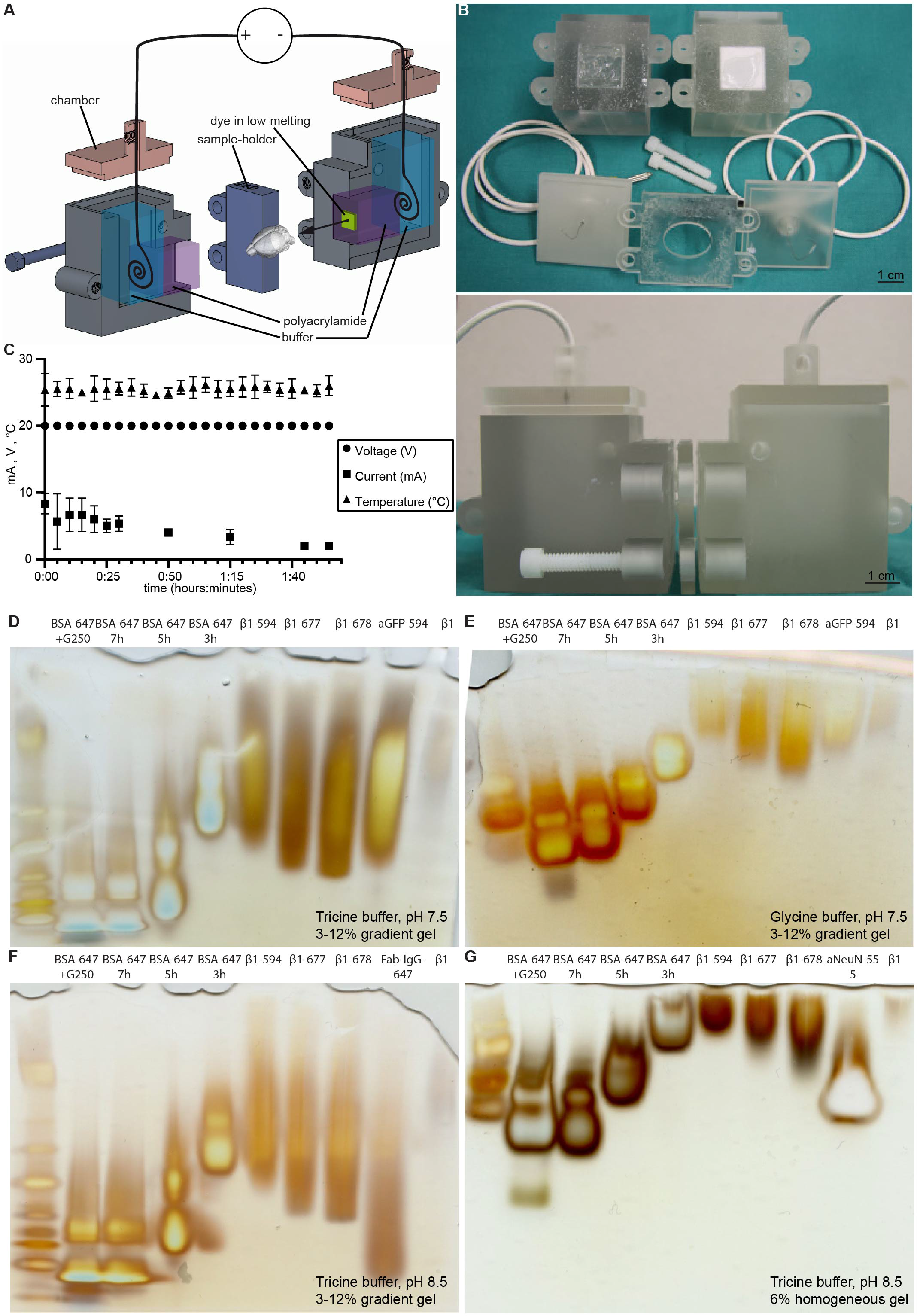
Digitally designed and 3D printed setups for convenient electrophoretic staining of tissues. (**A**-**B**) Rendering and 3D prints of electrophoretic tissue staining Q3D chambers. Electrophoresis buffer (light blue) chambers (gray) are separated by two blocks containing polyacrylamide plugs (purple). The sample is placed in a spacer (dark blue) between the plugs. One of the plugs contains the dye solution immobilized in low-melting agarose (green). Each chamber contains a platinum electrode (black). (**C**) Electrophoretic staining was performed at constant voltage and tissue temperature. The electrical current decreased after some minutes. (**D**-**G**) Non-denaturing polyacrylamide gel electrophoresis of various proteins at a range of pH and buffer compositions. (**D**-**F)** Tricine buffer provided better electrophoretic mobility than (E) glycine buffer, as did higher pH (<u>D</u>: pH 7.5, F: pH 8.5). (**D**-**G**) Increasing the electric charge of the β1-antibody by covalently coupling three valent anionic fluorophores (β1-594, β1-677, β1-768) increased electrophoretic mobility. BSA-647 was loaded in 2-hour intervals. In a 6% polyacrylamide native gel (G, while D - F are 3-12 % gradient gels) this resulted in running distances proportional to the run-time. (BSA-bovine serum albumin, β1 - β1 antibody, aNeuN-555-Alexa555 conjugated anti-NeuN antibody, aGFP-594-Alexa594 conjugated anti-GFP antibody).

**Supplementary Figure 3.**
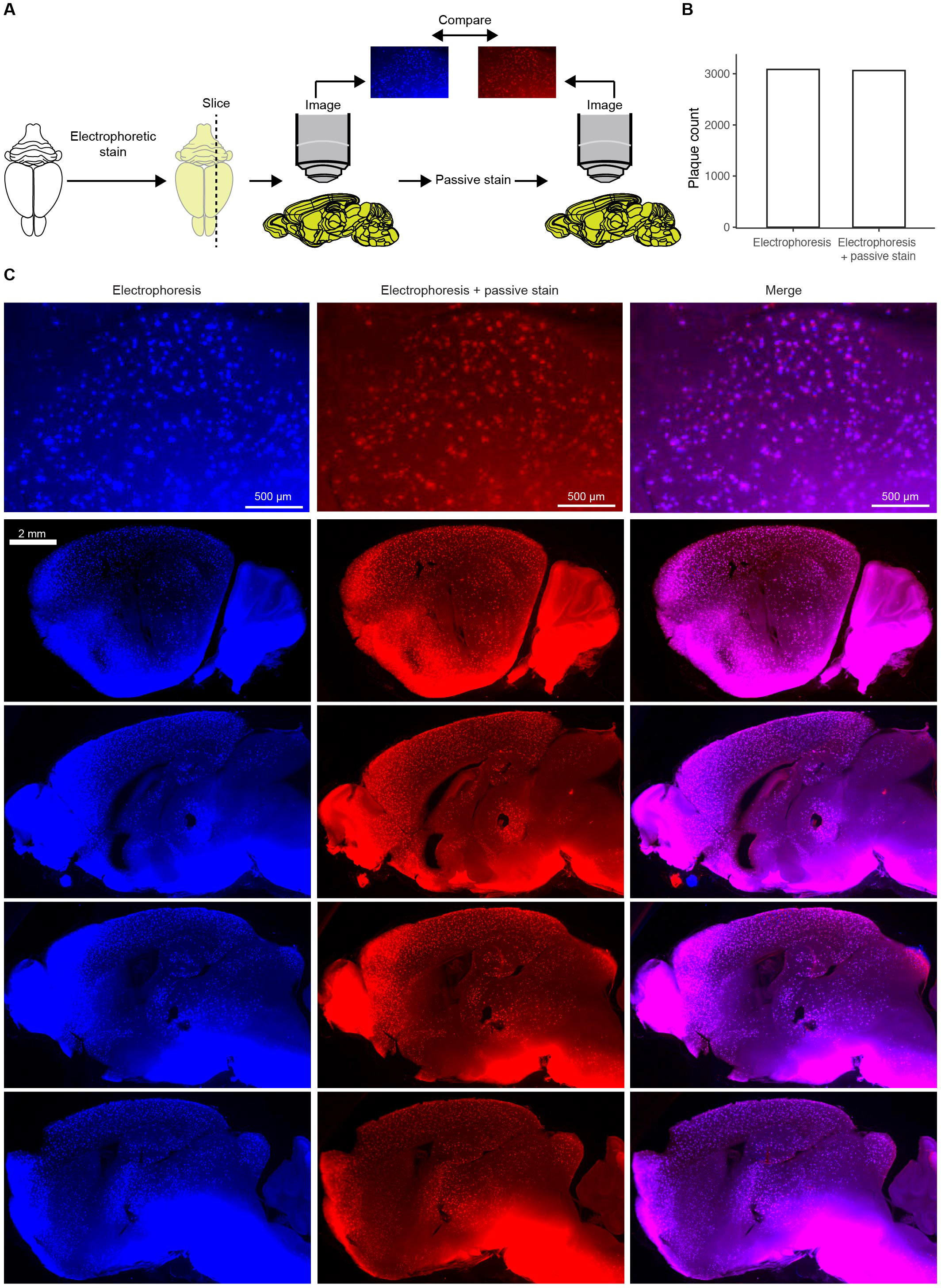
Identical Aβ plaque populations are labeled after electrophoretic and passive polythiophene staining. **(A) A clarified** APP/PS1 mouse brain was electrophoretically stained with qFTAA and hFTAA. Sagittal 500-μm sections were imaged with a fluorescent stereomicroscope. After the first round of imaging the slices were passively stained with qFTAA and hFTAA, followed by re-imaging. The images from the two imaging round were compared. (**B**) No significant difference between the number of plaques in electrophoretically stained and passively re-stained slices. (**C**) Visual inspection of the electrophoretically stained and passively re-stained slices shows close to identical plaque populations. Slight differences between the images are most likely due to the physical manipulations during re-staining and re-imaging.

**Supplementary Figure 4.**
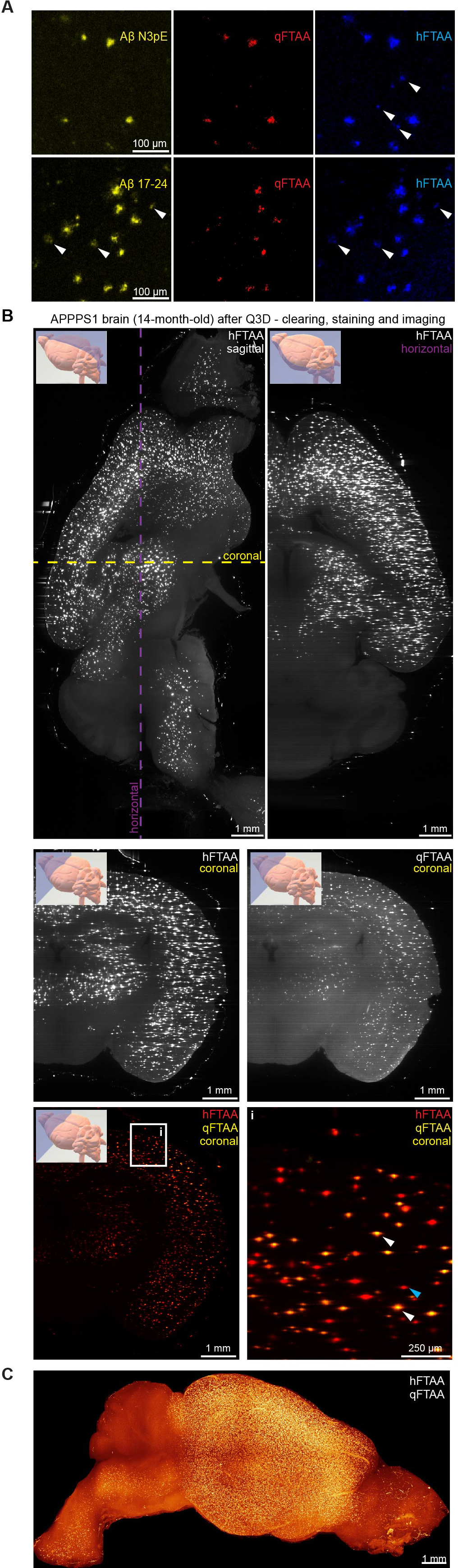
Homogeneous whole-brain Aβ plaque stain by electrophoretic infusion of qFTAA and hFTAA polythiophenes. (**A**) Plaque maturation states determined with qFTAA vs hFTAA were compared to those identified by N3pe (pyroglutamylated Aβ) vs Aβ_17-24_ (all Aβ moieties). APPPS1 brain sections (paraffin, 10 μm) were stained with Aβ-N3pE or Aβ17 24 followed by qFTAA+hFTAA staining. Aβ-N3pE detected plaques similarly to qFTAA, while hFTAA highlighted additional plaques. Conversely, Aβ_17-24_ labeled the entire plaque population similarly to hFTAA (white arrowheads). (**B**) Lightsheet imaging and digital reslicing of cleared whole brains. Plaques were visible in the cortex and in deep diencephalic areas, indicative of homogeneous dye penetration and image acquisition. Plaque maturity was assessed with qFTAA+hFTAA co-staining. As an example, hFTAA identified all plaques (blue arrows) whereas qFTAA stained the cores (white arrows) of more mature plaques. (**C**) Whole brain hemisphere rendering of an APPPS1 mouse with hFTAA signal. The cerebellum, where the Thy1 promoter is inactive, was unaffected.

**Supplementary Figure 5.**
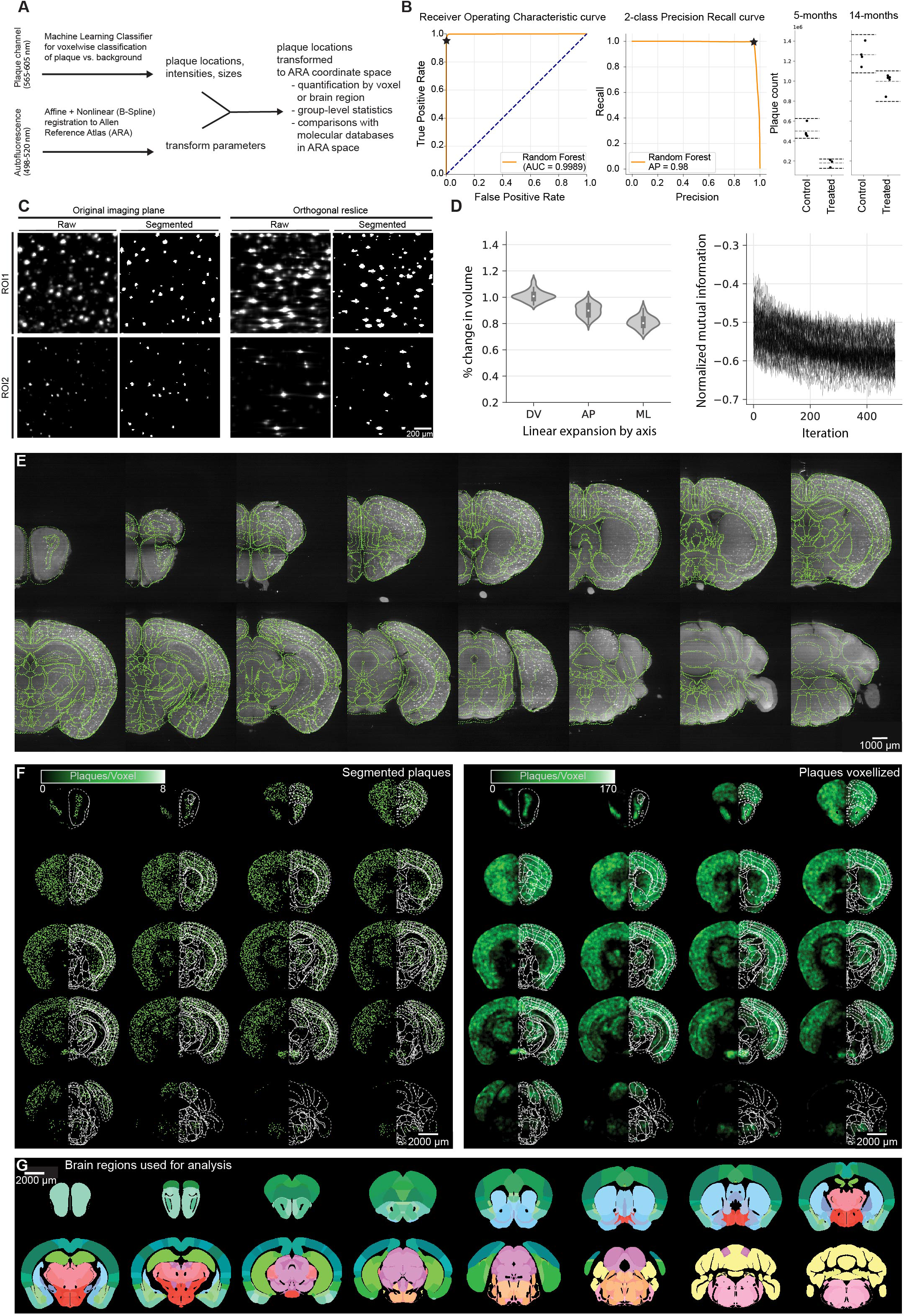
Digital atlas registration and machine learning-based plaque segmentation results in precise plaque-load heatmaps. (**A**) Schematic of sample quantification pipeline. Steps in the data analysis pipeline include accurate registration of brains to an atlas using non-linear transformations, plaque segmentation with expert-trained machine learning., and transformation of plaque information to an atlas space where group-level and other analyses can be performed. (**B**) The Receiver Operating Characteristic and Precision Recall curves quantify the classifier’s accuracy of plaque detection on an expert-annotated test dataset. When accounting for the classifier’s worst-case recall rate (dotted lines, 0.95%), the effect on whole brain plaque count following the NB360 treatment is greater than the variability from overcounting or undercounting whole brain plaques. (**C**) Machine learning-based segmentation fits with raw data. Spherical plaques (Orthogonal imaging plane) show spindle artefacts when digitally resliced orthogonally (Orthogonal reslice – Raw). Segmentation decreased artefacts (Orthogonal reslice – Raw and Segmented). (D) Singular values from the registration process capture linear expansion of cleared brains across three spatial axes with respect to the brain atlas. There was consistent shrinking across the anterior-posterior (AP) and medial-lateral (ML) axes, while the dorsal-ventral (ML) axis showed minimal linear expansion. Despite linear expansion, registration optimization metrics all converged to suitable values and (**E**) atlas region contours (green outlines) accurately align with the autofluorescence channels in three dimensions. (**F**) Segmented plaques (left) are voxelized resulting in heatmaps of either plaque density, mean size or maturity (right). (**G**) Statistics on segmented plaques can be done by grouping them into brain regions.

**Supplementary Figure 6.**
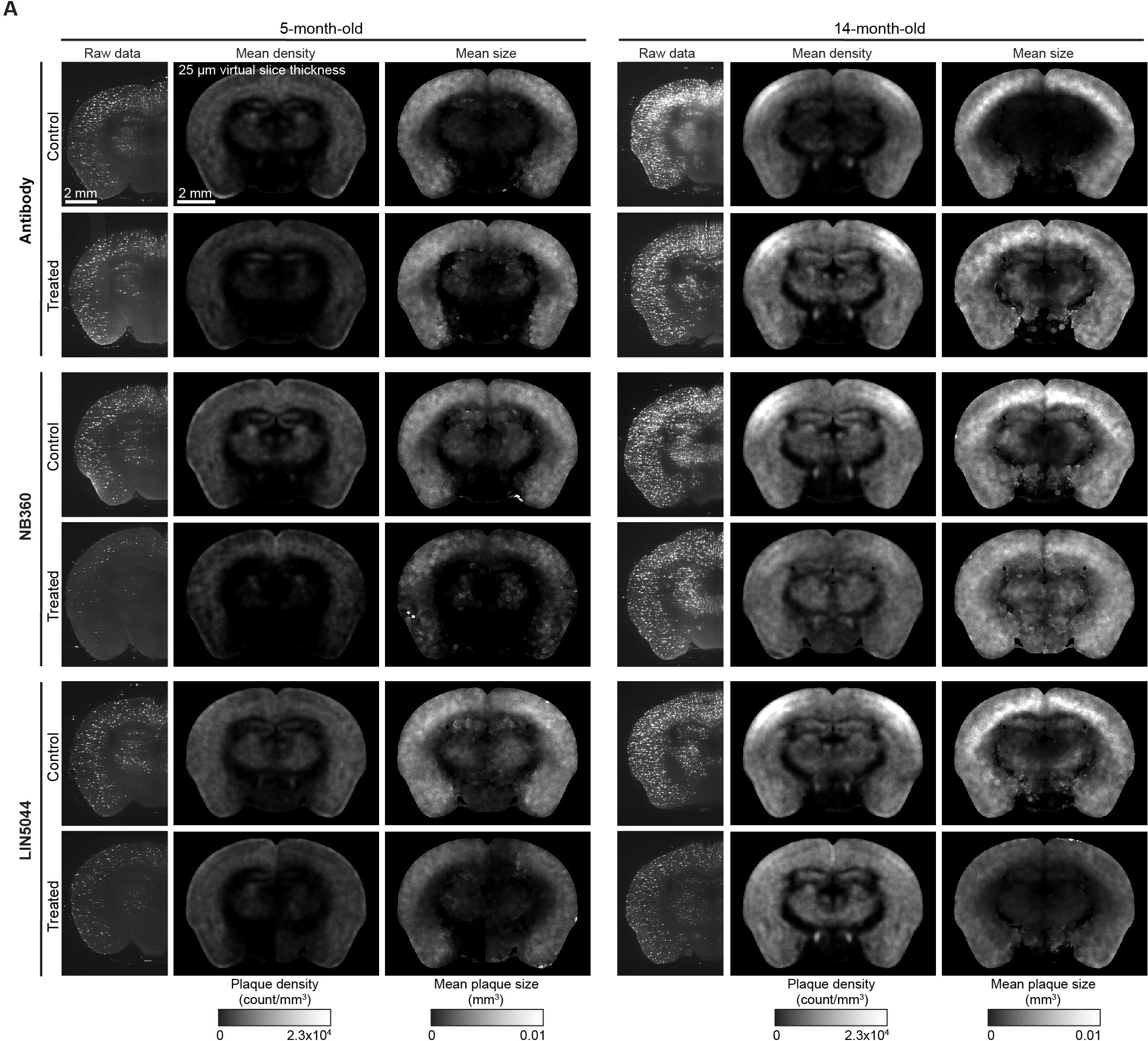
Data representations indicate differences between brains of control and treatment cohorts. (**A**) Heatmaps of mean plaque densities and sizes are shown across all the treatment and control groups. For each cohort an optical brain slice from lightsheet data is shown (Raw data). Plaque loads in aged mice are much higher than in young mice. The BACE1 inhibitor NB360 treatment suggests reduced plaque counts in 5 – month – old mice. LIN5044 treatment suggests decreased mean plaque size in 14 – month – old mice. In some cases, the optical brain slices do not precisely represent the anatomy of the atlas-registered heatmap-slices. This is because the mounting (and thus the orientation) of the brains in the microscope is variable.

**Supplementary Figure 7.**
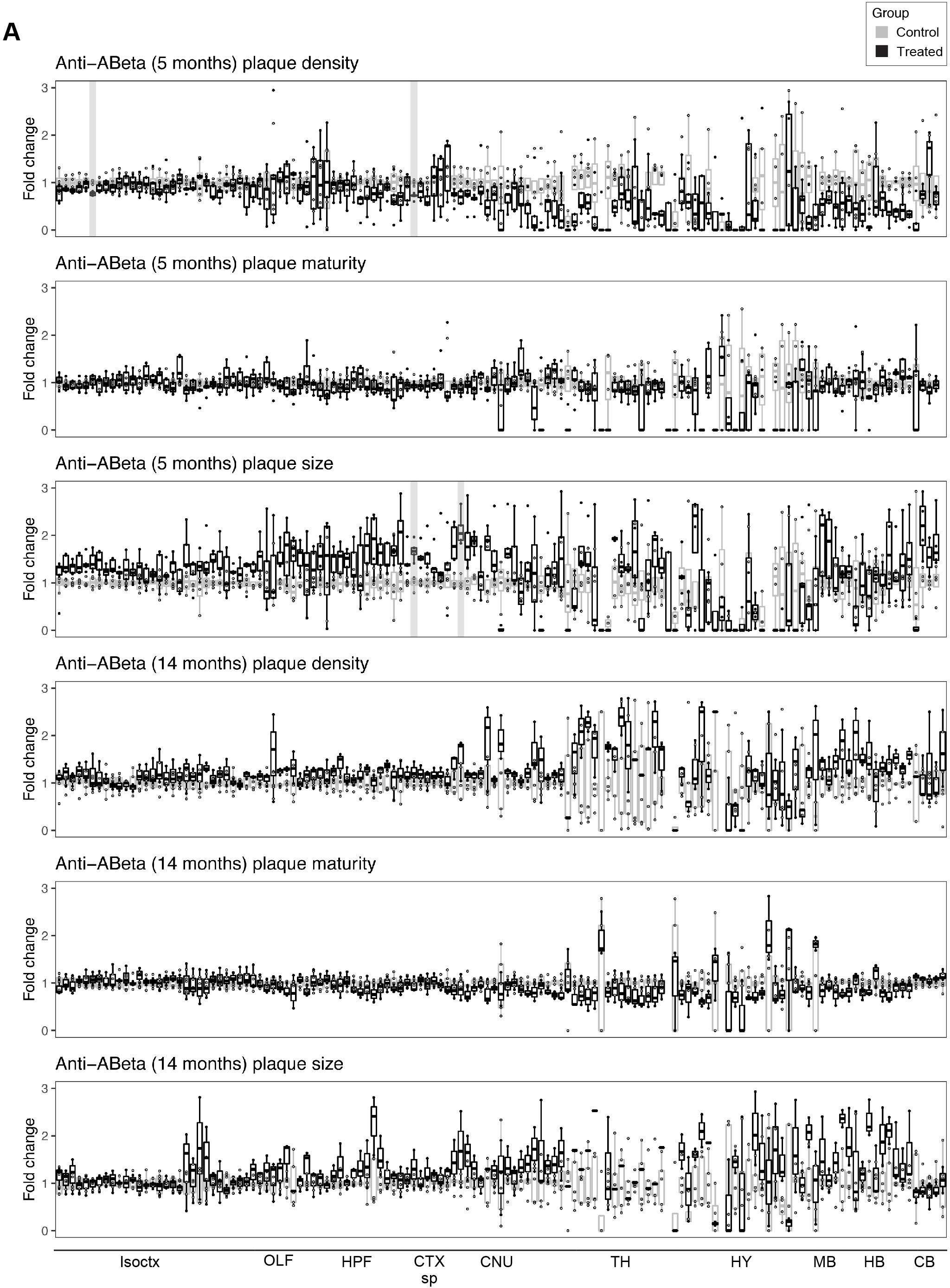
The β1 antibody has minimal to no effect on neuroanatomical plaque count, maturity, or size in 5-month-old and 14-month-old mice. (**A**) Fold-change reduction in various plaque metrics across all brain regions in both 5-month-old and 14-month-old mice, compared to control. The β1 antibody has a minimal effect on plaque-count increase in 5-month-old mice and has no effect in old mice. In 5-month-old mice, significant effects in plaque size reduction occurred mostly in the brainstem. The plaque-maturity change induced by β1 antibody treatment in 5- and 14-month-old mice is very limited. The plaque-size change induced by β1 antibody treatment in 5-month-old mice is very limited and absent in 14-month-old mice. The plaque-maturity change induced by β1 antibody treatment in 5- and 14-month-old mice is very limited. Brain regions with a significant treatment effect (p < 0.05) are shaded gray. Isoctx – isocortex, OLF – olfactory areas, HPF – hippocampal formation, CTX sp – cortical subplate, CNU – caudate nucleus, TH – thalamus, HY – hypothalamus, MB – midbrain, HB – hindbrain, CB – cerebellum.

**Supplementary Figure 8.**
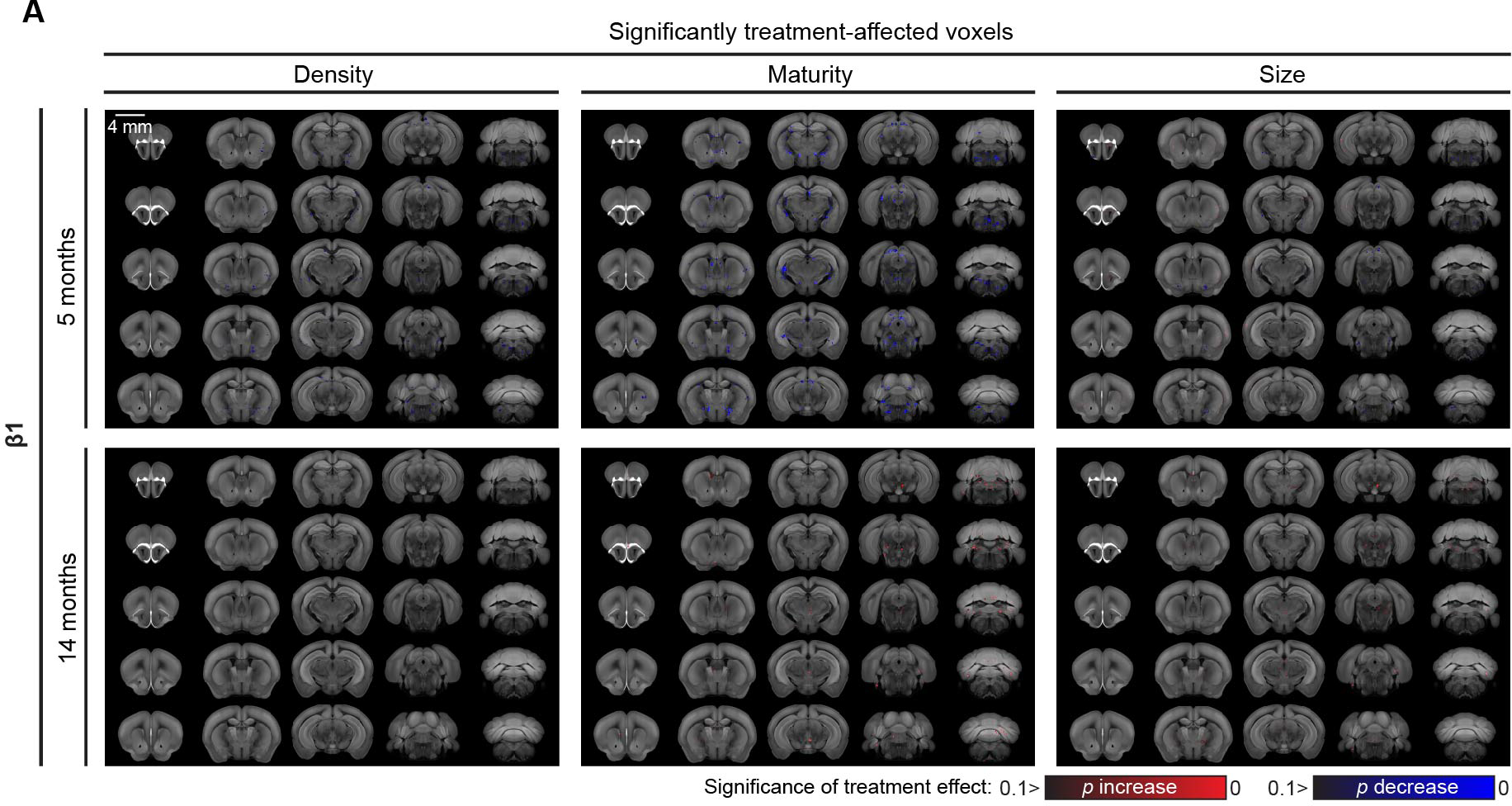
The β1 antibody has minimal to no effect on individual voxel plaque count, maturity, or size in 5-month-old and 14-month-old mice. (**A**) Heatmap of voxel-based statistics with significant mean plaque size reduction upon β1 antibody treatment in 5-month-old and 14-month-old mice, compared to control. The β1 antibody treatment shows a limited effect across all analyzed metrics.

**Supplementary Figure 9.**
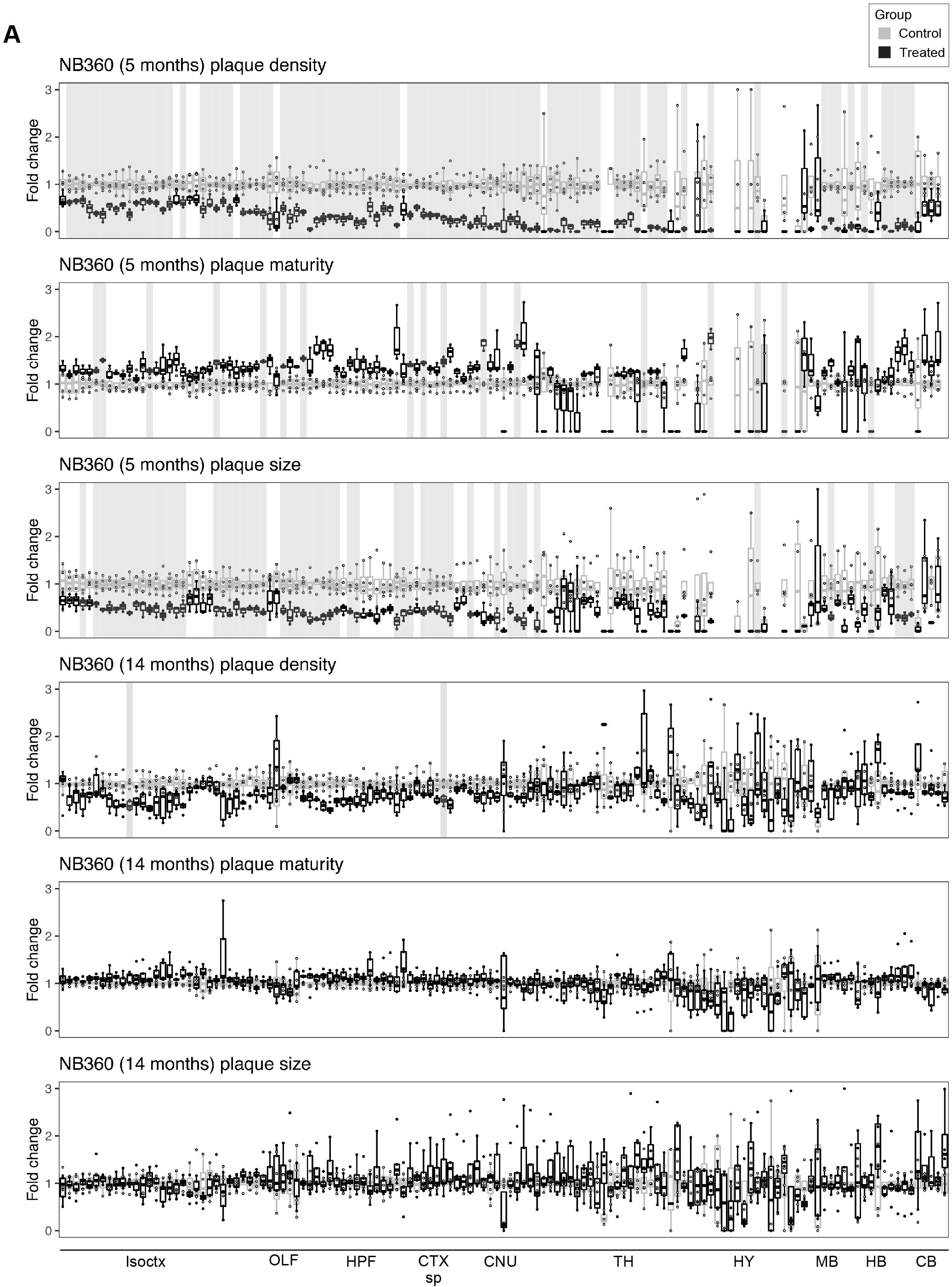
BACE1-inhibition induced conspicuous plaque count reduction in 5-month-old mice, but not in 14-month-old mice. (**A**) Heatmap of voxels with significant plaque count reduction upon NB360 treatment in young mice, compared to control. Upon BACE1 inhibition in 14-month-old mice, only very few voxels show significant plaque-count reduction. BACE1-inhibition induces considerable plaque size reduction in 5-month-old mice, but not in 14-month-old mice. Heatmaps of significantly affected voxels show very limited plaque size change by BACE1-inhibition in 14-month-old mice. Plaque maturity change by BACE1-inhibition in 5-month-old mice shows region-dependent maturity increase and decrease while there is no effect at 14-months. Brain regions with a significant treatment effect (p < 0.05) are shaded gray. Isoctx – isocortex, OLF – olfactory areas, HPF – hippocampal formation, CTX sp – cortical subplate, CNU – caudate nucleus, TH – thalamus, HY – hypothalamus, MB – midbrain, HB – hindbrain, CB – cerebellum.

**Supplementary Figure 10.**
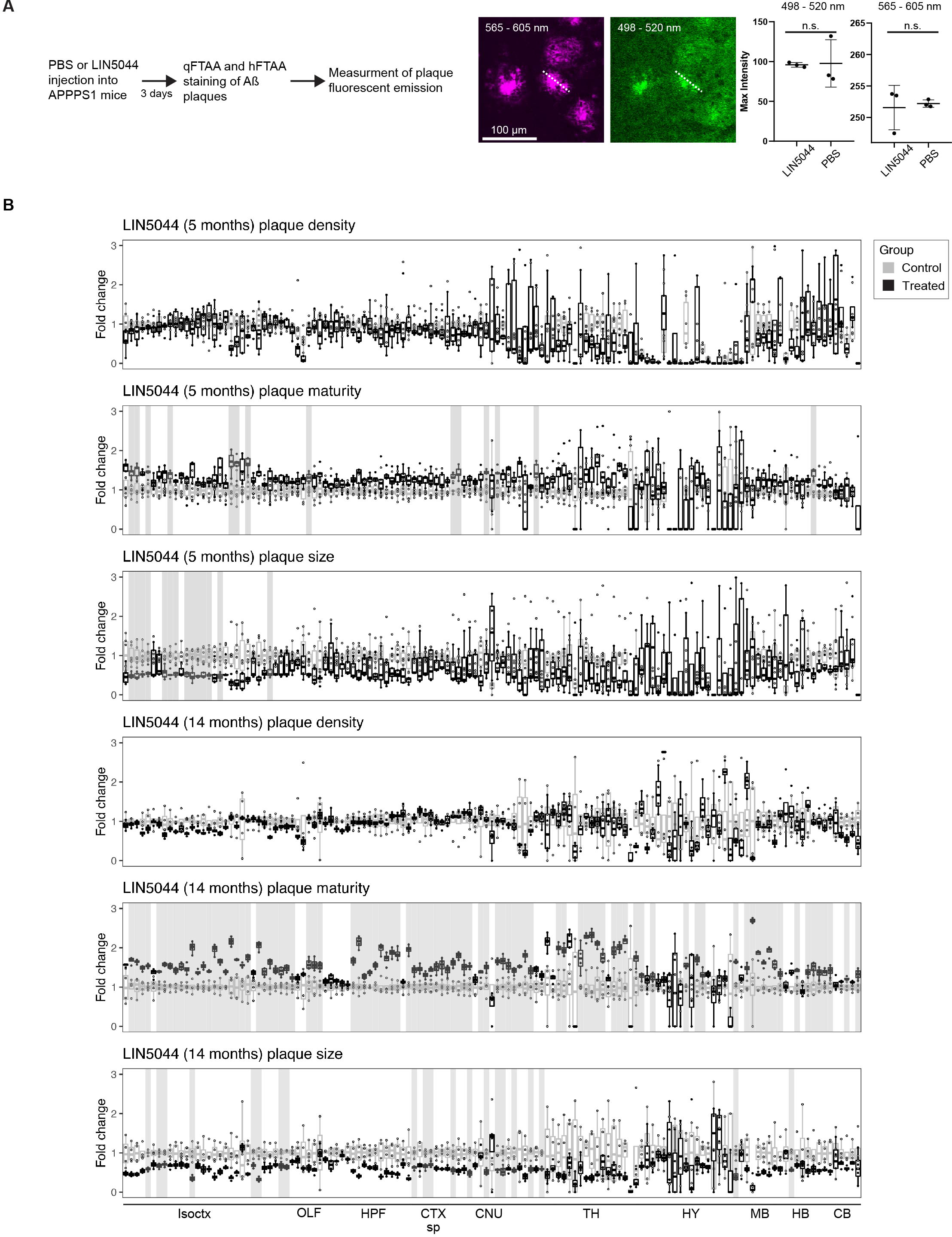
Trend in plaque-count reduction in few anatomical regions after LIN5044 treatment in 5- and 14-month-old mice. (**A**) To see whether LIN5044 induces artefacts into the plaque-maturity analysis APPPS1 mice were injected with a single-dose PBS (n=1) or LIN5044 (0.4 mg in 100μl PBS) (n=1). Next, emission intensities used for plaque maturity analysis (at wave lengths 498-520 and 565-605 nm) were measured in 10 plaques in 3 slices per animal with a confocal microscope. Plaques of LIN5044 and PBS injected mice showed no difference in fluorescent emissions. (**B**) Fold-change reduction in various plaque metrics across all brain regions in both 5-month-old and 14-month-old mice, compared to control. Plaque-count reduction is not significant but shows a trend in 14-month-old mice. Mean plaque-sizes are significantly reduced in some anatomical regions after LIN5044 treatment in 5-month-old mice, and in cortical areas of 14-month-old mice. Plaque maturity change by LIN5044 in 5-month-old mice shows some neuroanatomical regions with mean maturity increase, and a more widespread regional increase at 14-months in cortical areas. Brain regions with a significant treatment effect (p < 0.05) are shaded gray. Isoctx – isocortex, OLF – olfactory areas, HPF – hippocampal formation, CTX sp – cortical subplate, CNU – caudate nucleus, TH – thalamus, HY – hypothalamus, MB – midbrain, HB – hindbrain, CB – cerebellum.

**Supplementary Figure 11.**
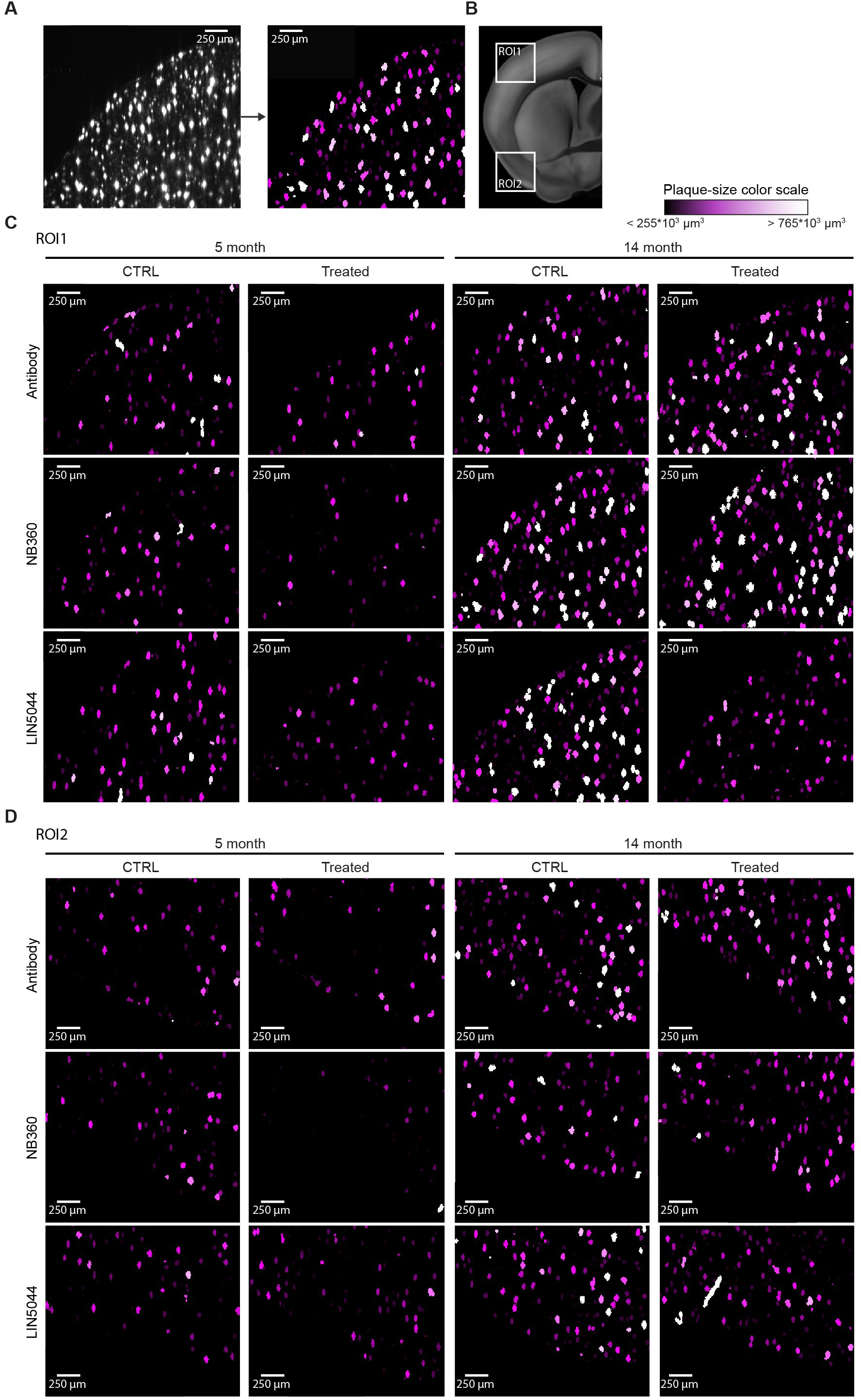
Raw data shows differential effects on plaque counts and sizes after amyloid β treatments. (**A**) We randomly picked an optical slice from the cortex (**B**, ROI1, ROI2 white inserts) of a control and a treated brain in each treatment cohort. Next, we segmented the plaques and color-coded them such that colors represent 4 equal portions of the total range of plaque sizes. (**C-D**) NB360 conspicuously reduced numbers and sizes of plaques in young mice, while LIN5044 the number of large plaques in aged mice. The effect in the other treatment cohorts is less obvious.

**Supplementary Figure 12.**
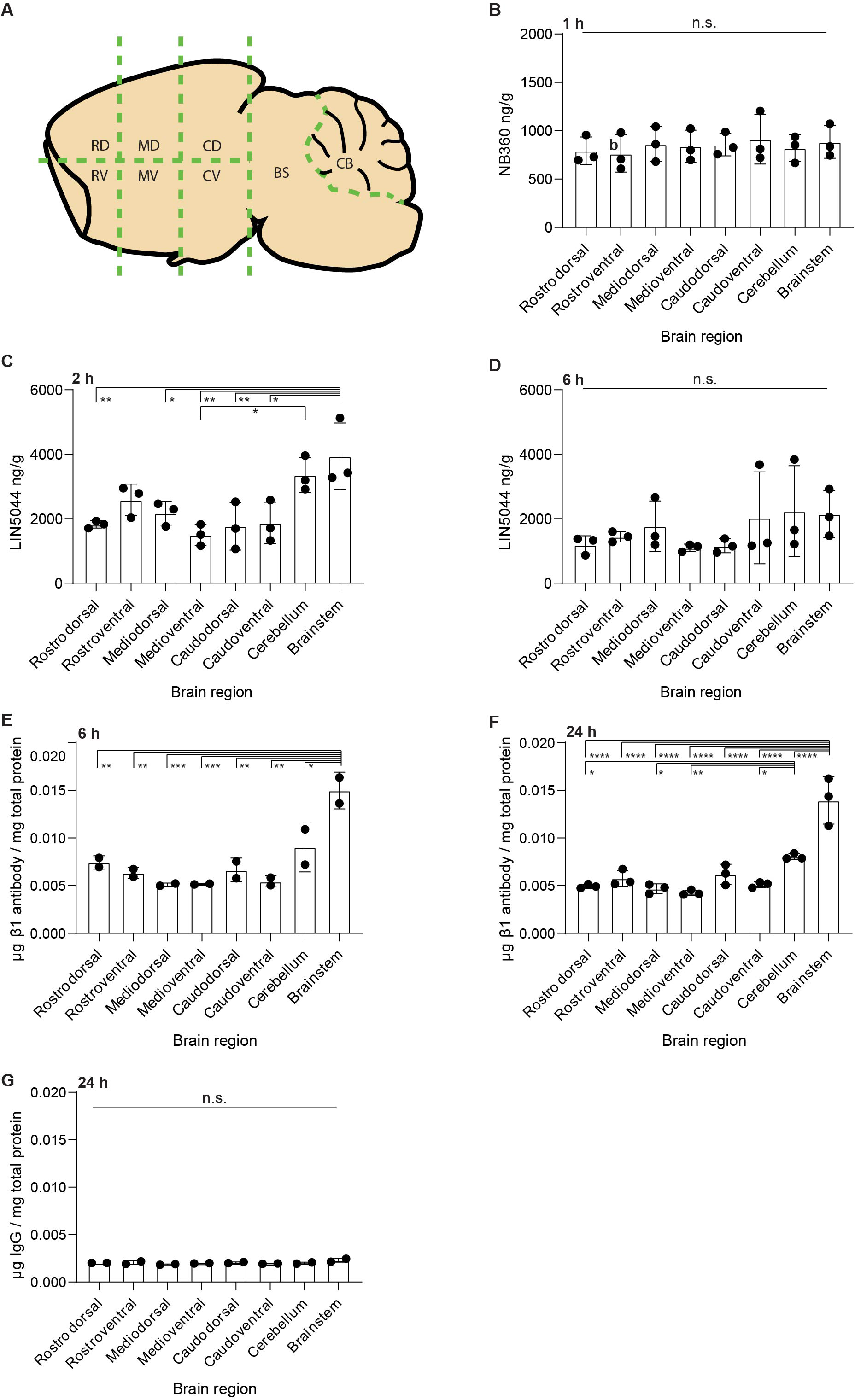
Variations in bioavailability across bulk regions do not explain the spatial patterns of drug effectiveness. (**A**) Brains were dissected into eight bulk regions to test regional drug distributions. (**B**) No regional difference in NB360 drug levels detectable after oral administration, (**C**) while LIN5044 shows significantly higher levels in the brainstem (ranging 0.0023<p<0.0345) and the cerebellum (p = 0.0241) 2 hours after intraperitoneal injection, (**D**) but no difference detectable at 6 hours. (**E, F**) ELISA measurements of β1 antibody levels after intraperitoneal application showed higher levels in the brainstem (for multiple brain regions ranging p = 0.0223 to p<9×10^-4^ at 6 hours and p<10^-4^ at 24 hours) and the cerebellum (for multiple brain regions ranging 0.0072<p<0.034 at 24 hours) compared to other regions both at 6 and 24 hours-similarly to LIN5044. (**G**) Pooled non-specific recombinant IgG shows no difference in regional distribution. RD – rostrodorsal, RV – rostroventral, MD – mediodorsal, MV – medioventral, CD – caudodorsal, CV – caudoventral, CB – cereballum, BS – brainstem

**Supplementary Figure 13.**
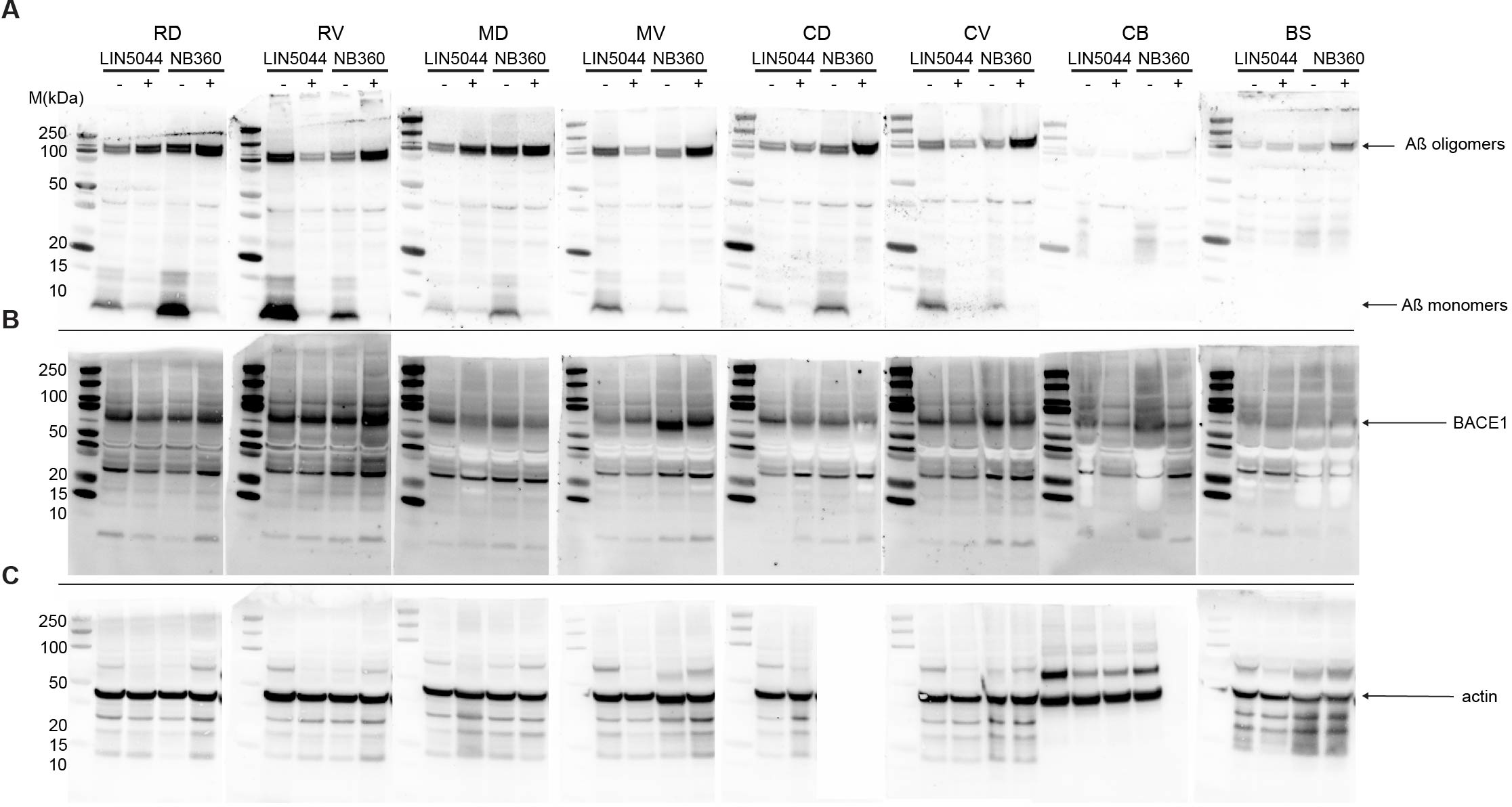
Biochemical measurements of Aβ and BACE1 levels across bulk regions do not explain spatial patterns of drug effectiveness. 2-month-old APPPS1 mice were treated with LIN5044 or NB360 for 3 months, followed by the dissection of one brain hemisphere into 8 bulk regions (**Figure S11A**). (**A**) Monomeric and oligomeric Aβ species are reduced by both LIN5044 and NB360 treatments in all 8 macroscopic regions when compared to control groups. NB360 has a stronger effect on monomers than LIN5044, while the opposite is true for oligomers. (**B**) BACE1 is detectable in all macroscopic regions without a clear regional predominance. Neither LIN5044 nor NB360 have a reproducible effect on BACE1 levels across macroscopic regions. (**C**) Actin is detectable in all macroscopic regions without a clear regional predominance. RD – rostrodorsal, RV – rostroventral, MD – mediodorsal, MV – medioventral, CD – caudodorsal, CV – caudoventral, CB – cereballum, BS – brainstem

**Supplementary Figure 14.**
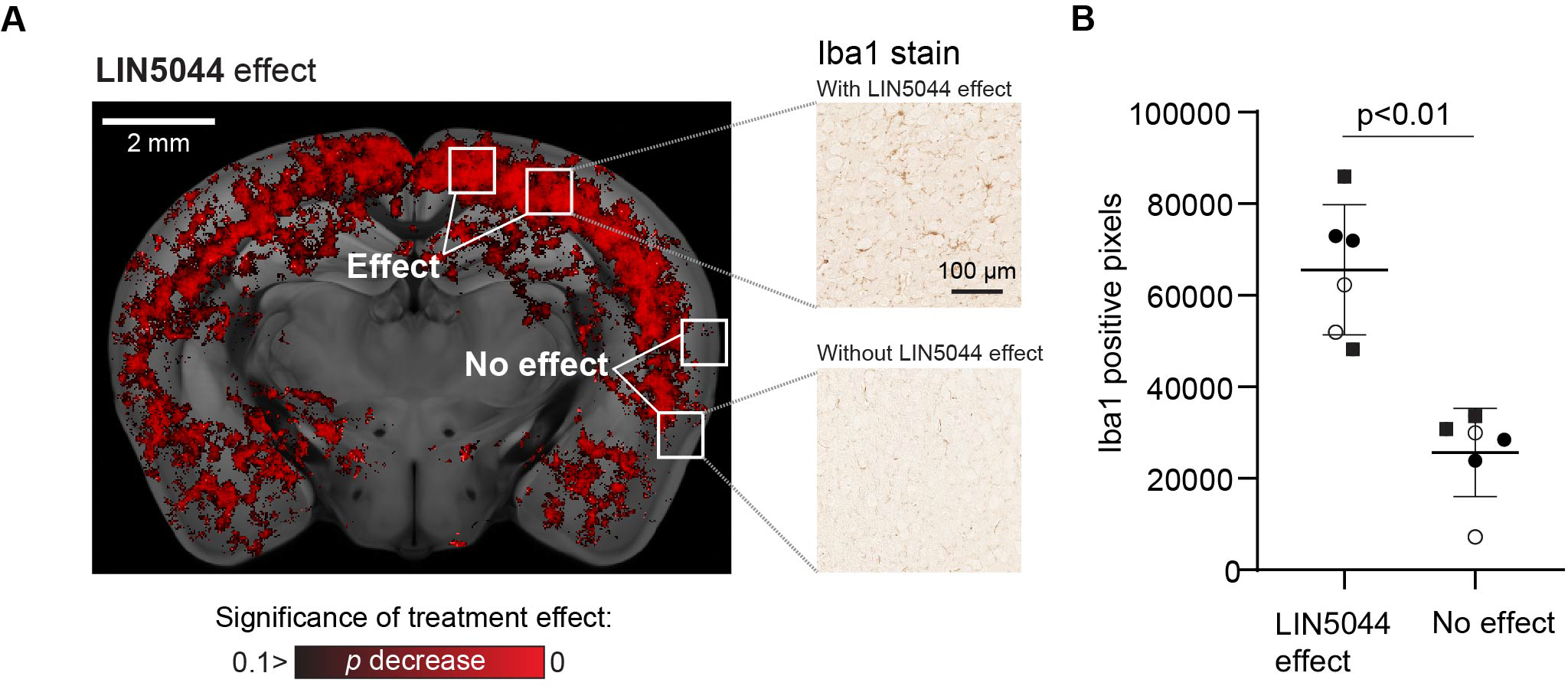
Histological sectioning and Iba1 labeling validates increased microglia density in regions where LIN5044 was more effective in reducing plaque size. (**A**) Regions of interest were selected based on voxel-based statistics of LIN5044 efficacy in reducing plaque size in 14-month-old mice. Microglia density was measured in regions displaying both strong or absent LIN5044 efficacy (3 three wild-type mice, 3 slices/mouse). (**B**) Regions showing strong LIN5044 effects contained more microglia (each distinct symbol represents one mouse).

**Supplementary Figure 15.**
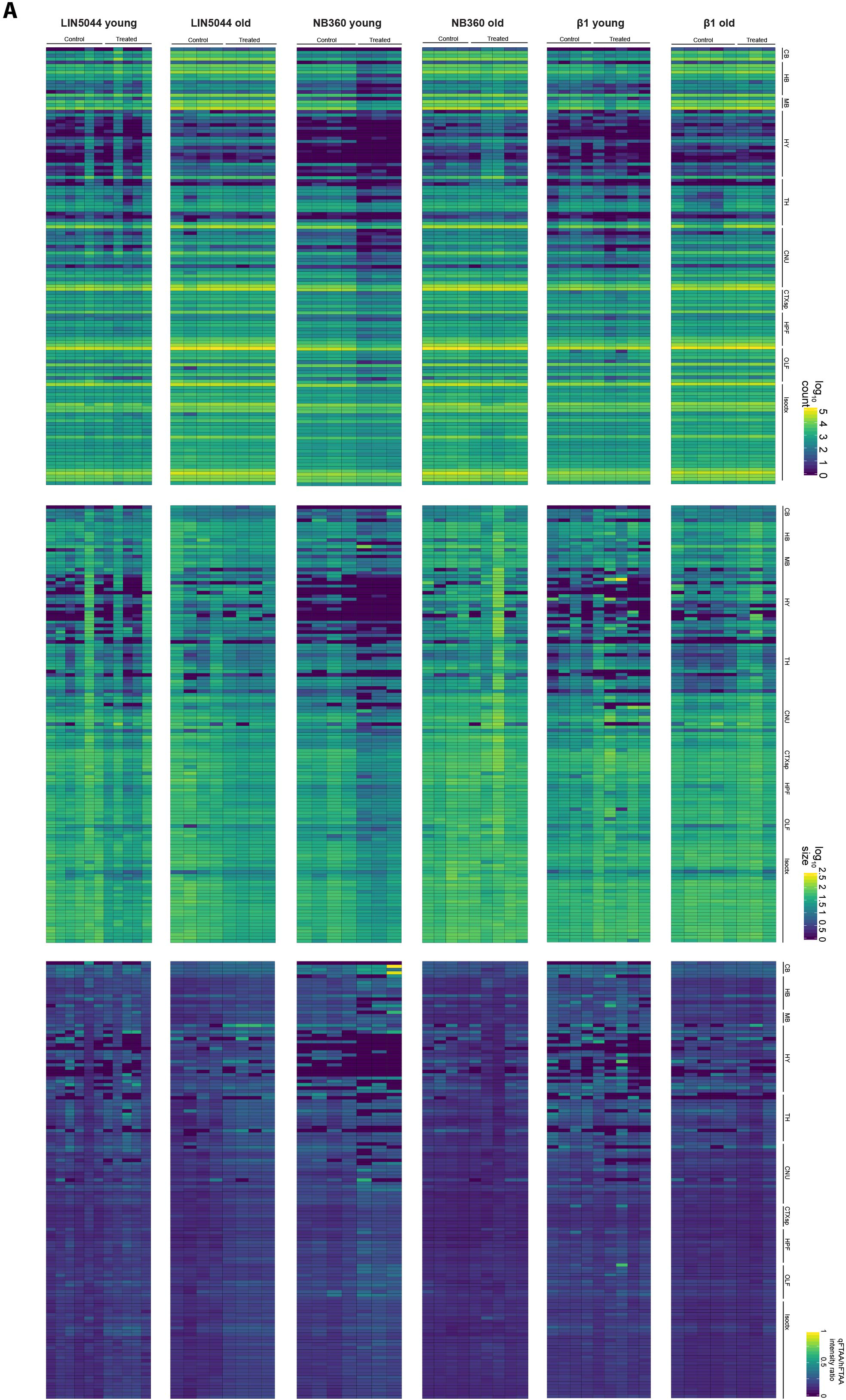
Per-subject regional quantification of plaque count, size, and maturity. (A) The regional quantification of total plaque count, mean plaque size, and mean plaque maturity for individual samples is depicted. Columns represent either a control or treated subject across the various cohorts. Regions are grouped by larger neuroanatomical structures: Isoctx – isocortex, OLF – olfactory areas, HPF – hippocampal formation, CTX sp – cortical subplate, CNU – caudate nucleus, TH – thalamus, HY – hypothalamus, MB – midbrain, HB – hindbrain, CB – cerebellum.

## Supplementary Tables

**Table S1.** Number of mice in treatment cohorts.

| | β1-old | β1-young | NB360-old | NB360-young | LIN504 4-old | LIN504 4-young |
| --- | --- | --- | --- | --- | --- | --- |
| control | 5 | 4 | 4 | 5 | 4 | 6 |
| treated | 3 | 6 | 5 | 3 | 4 | 6 |

**Table S2.**
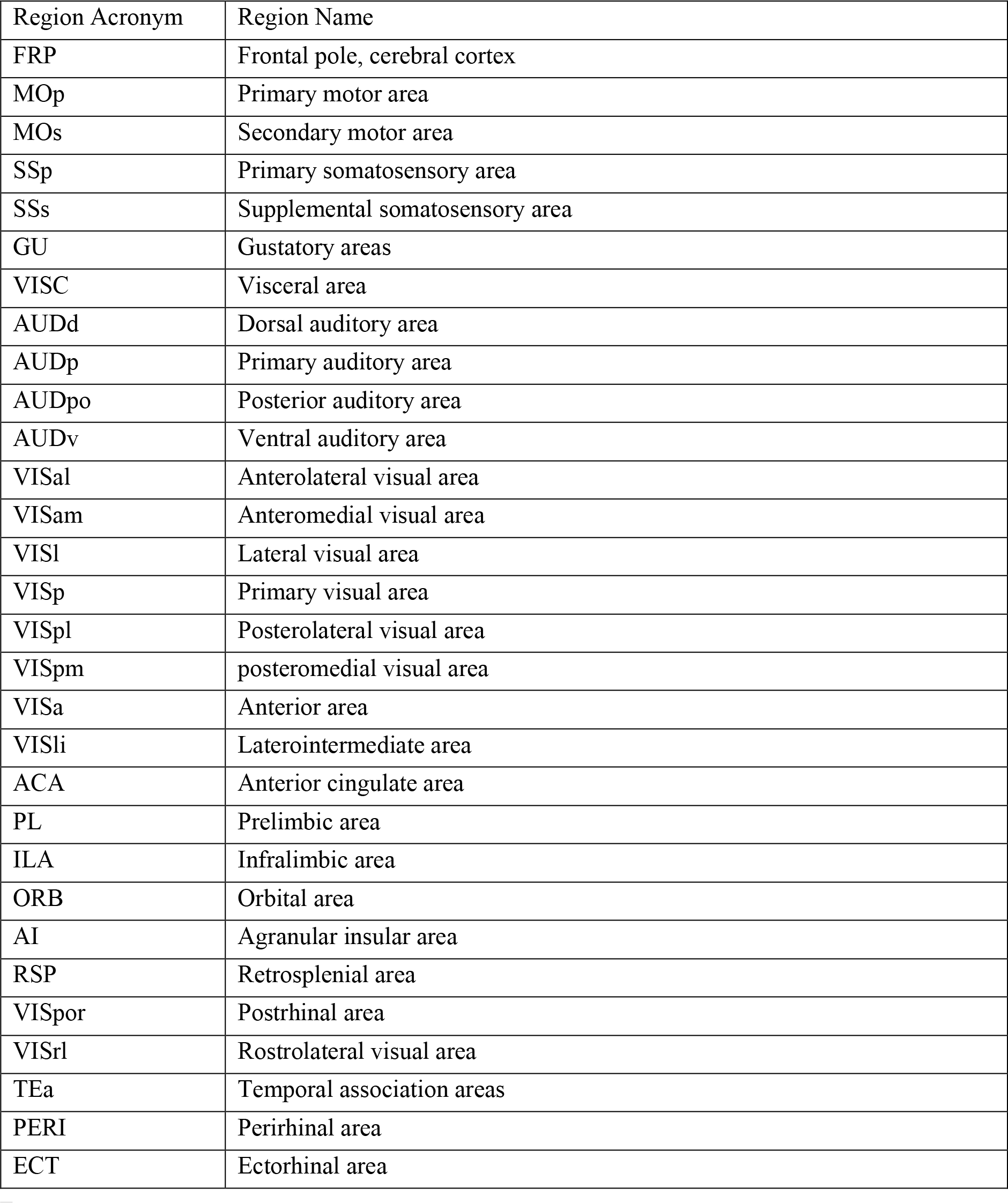
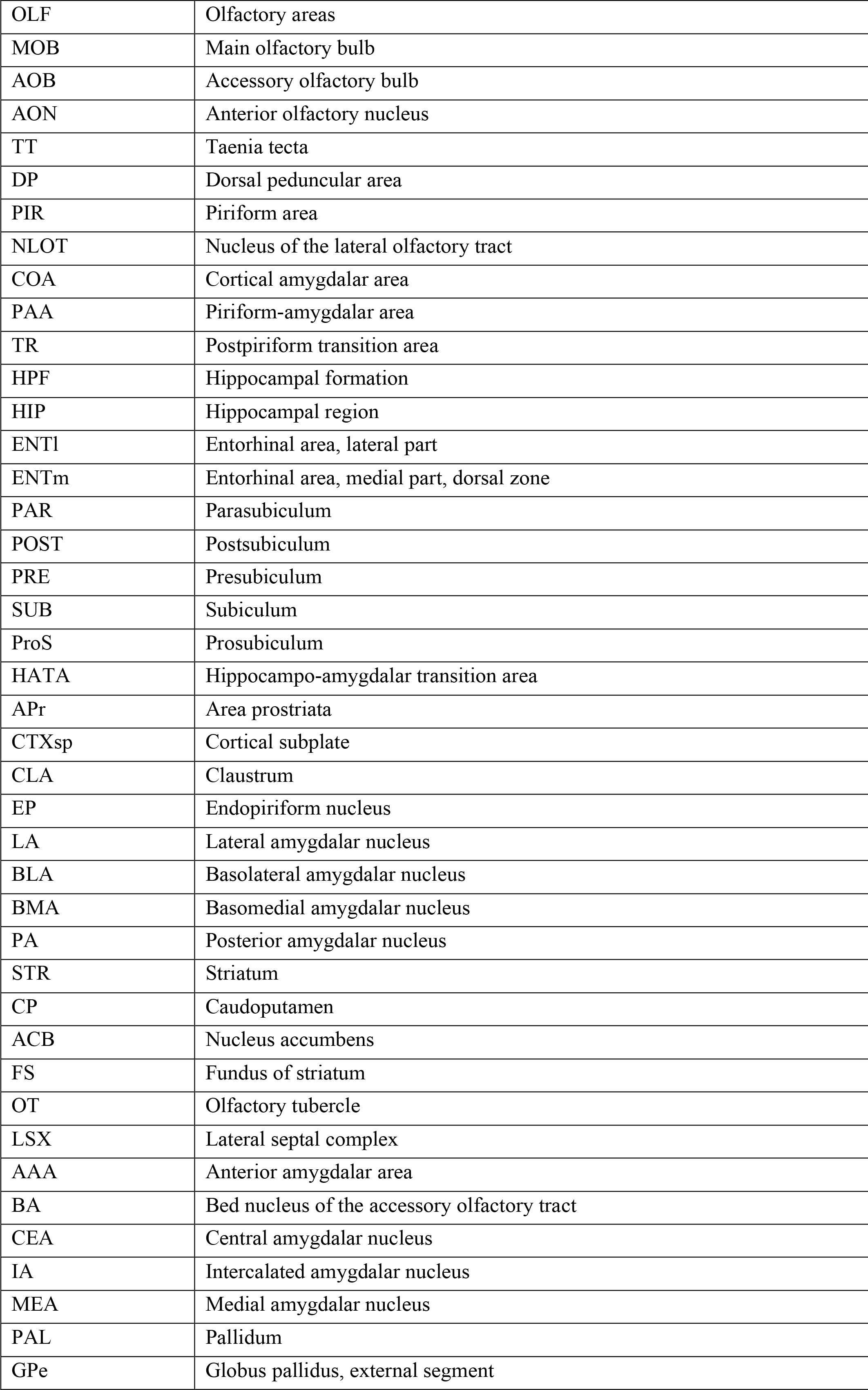
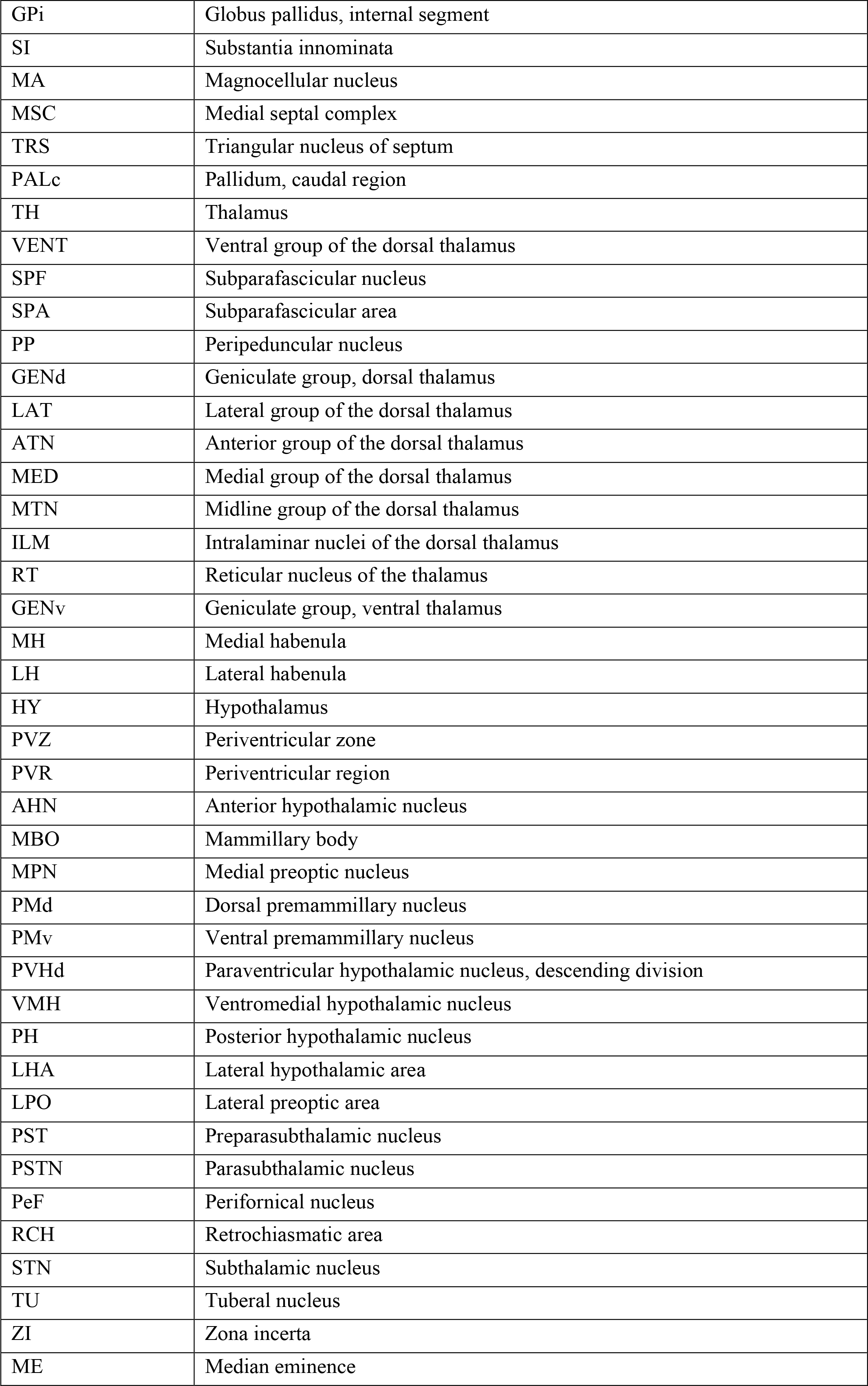
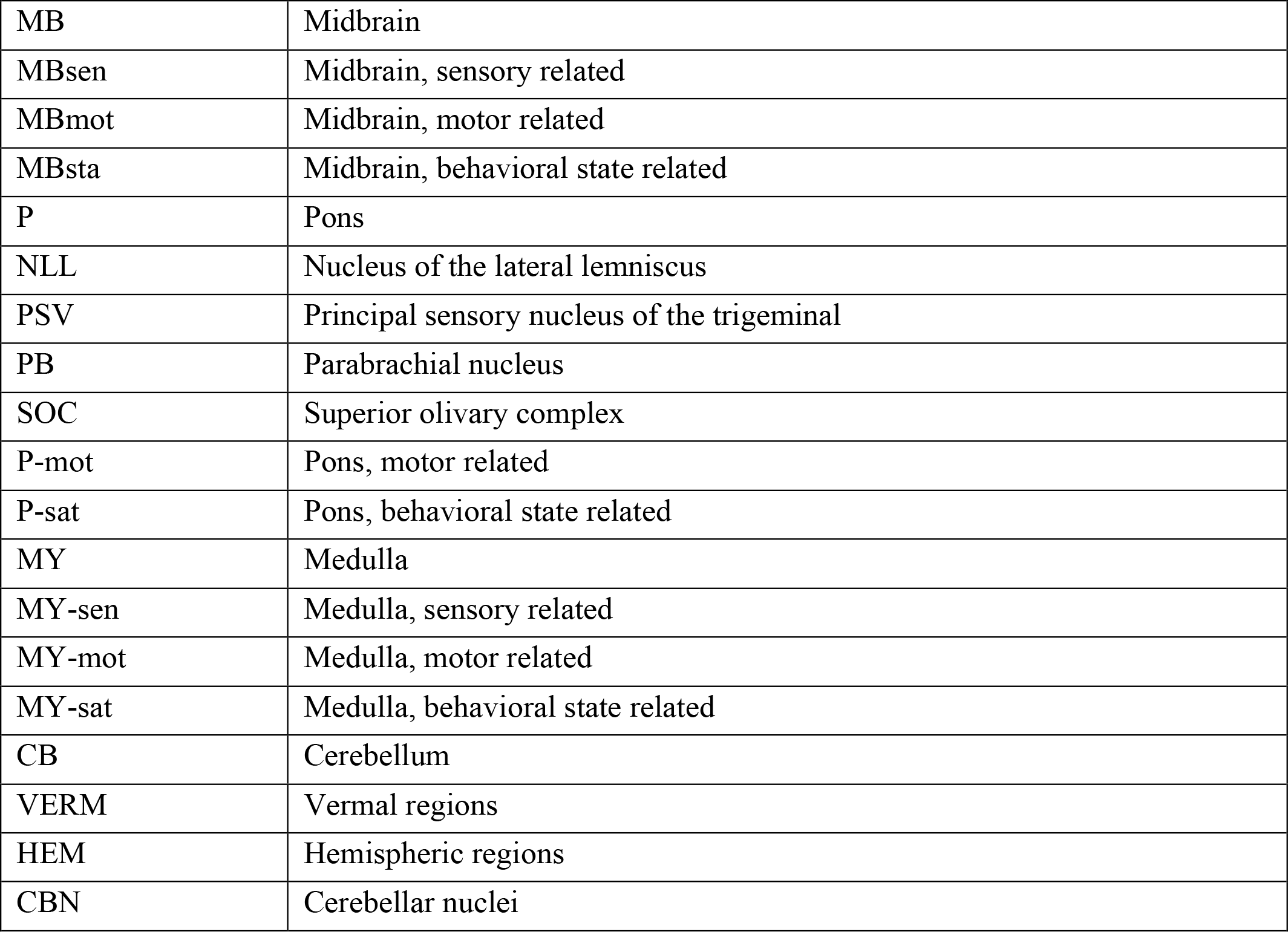
Neuroanatomical areas used for Allen Reference Atlas registration.

**Table S3.** Plaque loads in control mice of each treatment cohort show low variability.

| Treatment | Age | Count Mean | Count SD | Count SE |
| --- | --- | --- | --- | --- |
| AB | old | 2108735 | 292186 | 130670 |
| AB | young | 1097711 | 159214 | 79607 |
| BACE1 | old | 2790290 | 245900 | 122950 |
| BACE1 | young | 1151097 | 159849 | 79924 |
| LCP | old | 2738744 | 333892 | 166946 |
| LCP | young | 1243608 | 156807 | 64016 |
| <all> | old | 2512293 | 426799 | 118373 |
| <all> | young | 1175491 | 159340 | 42585 |

**Table S4.** Significantly affected voxels (SAV) after NB360, β1-antibody or LIN5044 treatment. After all the brain scans were registered to a brain atlas, the brains were deconstructed into standard voxels in a coordinate system. Then, for every treatment a “phantom” brain-volume was generated where each voxel represented the p-value of treatment effect. This was generated by two-sided t-testing all treated against all control brains (for a respective voxel). Voxels with p-values either p<0.1 or 0.05 were termed as SAV. Each treatment’s phantom brain was thresholded to only contain SAVs (p<0.1 or 0.05). The effect-overlap between treatments was defined by the voxels which were significantly affected in both of the compared treatments. Our results show that most SAVs are non-overlapping. The overlap was <1% or <2.65% at p<0.05 or p<0.1, respectively.

| $p<0.05$ | | | | | | | |
| --- | --- | --- | --- | --- | --- | --- | --- |
| Young |  |  |  | Old |  |  |  |
| Density | NB360 | $\beta 1$ | LIN5044 | Density | NB360 | $\beta 1$ | LIN5044 |
| NB360 | 2'649'191 | 435 | 2 | NB360 | 46 | 0 | 0 |
| $\beta 1$ | 435 | 4'625 | 0 | $\beta 1$ | 0 | 40 | 0 |
| LIN5044 | 2 | 0 | 29 | LIN5044 | 0 | 0 | 276 |
| Size | NB360 | $\beta 1$ | LIN5044 | Size | NB360 | $\beta 1$ | LIN5044 |
| NB360 | 66'128 | 3 | 1 | NB360 | 10 | 0 | 0 |
| $\beta 1$ | 3 | 2'518 | 0 | $\beta 1$ | 0 | 18 | 1 |
| LIN5044 | 1 | 0 | 80 | LIN5044 | 0 | 1 | 492'779 |
| Maturity | NB360 | $\beta 1$ | | Maturity | NB360 | $\beta 1$ | |
| NB360 | 1'593'392 | 9'542 |  | NB360 | 136 | 0 |  |
| $\beta 1$ | 9'542 | 52'893 | | $\beta 1$ | 0 | 639 | |
| $p<0.1$ | | | | | | | |
| Young |  |  |  | Old |  |  |  |
| Density | NB360 | $\beta 1$ | LIN5044 | Density | NB360 | $\beta 1$ | LIN5044 |
| NB360 | 7'185'540 | 5'443 | 30 | NB360 | 16304 | 0 | 0 |
| $\beta 1$ | 5'443 | 21'359 | 0 | $\beta 1$ | 0 | 69 | 0 |
| LIN5044 | 30 | 0 | 121 | LIN5044 | 0 | 0 | 486 |
| Size | NB360 | $\beta 1$ | LIN5044 | Size | NB360 | $\beta 1$ | LIN5044 |
| NB360 | 888'437 | 282 | 1'302 | NB360 | 25 | 0 | 0 |
| $\beta 1$ | 282 | 8'661 | 1 | $\beta 1$ | 0 | 61 | 5 |
| LIN5044 | 1'302 | 1 | 20'530 | LIN5044 | 0 | 5 | 2'941'884 |
| Maturity | NB360 | $\beta 1$ | | Maturity | NB360 | $\beta 1$ | |
| NB360 | 2'181'488 | 19'777 |  | NB360 | 221 | 0 |  |
| $\beta 1$ | 19'777 | 84'862 | | $\beta 1$ | 0 | 1432 | |

**Table S5.** Third-party libraries used for the computational pipeline.

| Software and Algorithms | Source |
| --- | --- |
| ClearMap | Renier, N. et al. Mapping of Brain Activity by Automated Volume Analysis of Immediate Early Genes. *Cell* **165**, 1789-1802, doi:10.1016/j.cell.2016.05.007 (2016). |
| Python | Python Software Foundation. Python Language Reference, version 2.7. Available at http://www.python.org |
| FIJI | Schindelin, J. et al. Fiji: An open-source platform for biological-image analysis. *Nat. Methods* **9**, 676–682 (2012). |
| R | R Core Team (2013). R: A language and environment for statistical computing. R Foundation for Statistical Computing, Vienna, Austria. http://www.R-project.org |

